# Complete Genome Sequence, Metabolic Profiling and Functional Studies reveal *Ligilactobacillus salivarius LS-ARS2* is a Promising Biofilm-forming Probiotic with Significant Antioxidant, Antibacterial, and Antibiofilm Potential

**DOI:** 10.1101/2024.12.04.626935

**Authors:** Sinjini Patra, Biswaranjan Pradhan, Anasuya Roychowdhury

## Abstract

**BACKGROUND:** Probiotics restore microbial balance and prevent gut-inflammation. Therefore, finding out novel probiotic strains is a demand. As gut-microbe, benefits of *Ligilactobacillus salivarius (LS)* are established. However, strain-specific detailed studies are limited. Here, we illustrate probiotic attributes of novel *LS-ARS2* for its potential application as food-supplement and/or therapeutic to improve gut-health.

**METHODS:** Whole genome sequencing (WGS) and phylogenetic analysis confirm the strain as *LS*. To establish probiotic properties, acid-bile tolerance, auto-aggregation, cell-surface-hydrophobicity, biofilm-formation and adhesion-assays are performed. To ensure safety attributes, antibiotic-susceptibility, haemolytic, DNase, trypan-blue and MTT assays are done. ABTS, DPPH, superoxide, hydroxyl free radical scavenging assays are used to determine anti-oxidant potential. Antibacterial assays including co-culture assay with pathogen and pathogenic biofilm-inhibition assays are performed to explore antibacterial efficacy. To characterize metabolic-profile of *LS-ARS2*-derived cell-free-supernatant (CFS), HRMS analysis are carried out. Consequently, WGS-analyses predict potential molecular associations related to functional outcomes.

**RESULTS:** We find *LS-ARS2* as remarkable fast-growing strain that shows acid and bile tolerance (>60% survival rate) indicating promising gut-sustainability. High auto-aggregation capacity (>80%), robust cell-surface hydrophobicity (>85%), and adhesion efficacy to Caco-2 cells illustrate significant potential of *LS-ARS2* for gut colonization. Fascinatingly, *LS-ARS2* is able to form biofilm within 24 h (p<0.0001), rare among *LS* strains, indicating the potential of the strain for efficient stay in the gut. The strain ensures safety attributes. *LS-ARS2*-WGS analysis recognizes probiotic-specific determinants, predicts genomic stability, identifies orthologous-clusters for diverse functions, and predicts metabolites and bacteriocins. HRMS-studies with *LS-ARS2-*CFS further validate the presence of diverse beneficial metabolites with antimicrobial and immunomodulatory potential. *LS-ARS2* shows significant antioxidant properties in ABTS (>60%), DPPH (>10U/mL), superoxide (>70%), and hydroxyl free radical scavenging assays (>70%). Further, *LS-ARS2* shows antimicrobial activities against Gram-positive Methicillin-resistant *Staphylococcus aureus (MRSA)* and Gram-negative multidrug-resistant clinical strains enterotoxigenic *Escherichia coli, Vibrio cholerae,* and *Shigella flexneri*. Anti-*Salmonella* effect of *LS-ARS2* is prominent (p<0.0001). Most interestingly, *LS-ARS2*-CFS inhibits *MRSA*-biofilm (p<0.0001), again rare among *LS* strains.

**CONCLUSIONS**

*LS-ARS2* is a novel fast-growing, biofilm-forming probiotic with significant antioxidant, antibacterial, and anti-biofilm potentials, suggesting the promising applications of *LS-ARS2* for combating pathogenic biofilms and improving gut-health. However, further *in vivo* studies would facilitate their potential applications.

## 1. Introduction

The human gut is colonized by 100 trillion bacteria along with fungi, viruses, protozoans, and archaea, which is collectively called gut-microbiota (Saxami et al., 2023). The gut microbiome is a dynamic ecosystem that manifests a symbiotic relationship with the host. It regulates host metabolism, and immunity and preserves intestinal barrier function by restoring a balance between beneficial and pathogenic bacteria in the gut. However, several factors, like diets rich in processed food and refined carbohydrates, unhealthy lifestyles, and uncontrolled medications, jeopardize the equilibrium of gut-microbial composition, which is termed dysbiosis (Pedroza Matute and Iyavoo, 2023). As a result, barrier integrity is disrupted and permeability is increased, leading to the entry of metabolites and endotoxins from pathogens. This triggers the activation of the immune cells and intestinal inflammation. Constant elevation of inflammatory mediators further promotes chronic inflammatory diseases like cancer, chronic respiratory diseases, atherosclerosis, stroke, and type 2 diabetes mellitus (Hakansson and Molin, 2011). As reported by the World Health Organization (WHO), inflammation-associated diseases are emerging as the most severe health hazard, with around 60% of all deaths across the world (Furman et al., 2019). Interestingly, probiotics can restore microbial balance in the intestine and reduce the severity of gut-inflammation (Patra et al., 2022; Patra et al., 2021). Therefore, probiotic intake as a diet or food supplement could be a preventive strategy for these inflammatory diseases.

Lactic acid bacteria (LAB) are the major source of probiotics. However, not all LAB are probiotics and should be evaluated for their probiotic attributes and safety profile (Ayed et al., 2024). According to the International Scientific Association for Probiotics and Prebiotics (ISAPP), for probiotic application, the strain must be tolerant to the harsh conditions (acidic and alkaline) to survive during passage through the gastrointestinal tract. It should have auto-aggregation and adhesion capacity to adhere and colonize the intestinal mucosa. After colonization, the bacteria must have beneficial effects, like the secretion of bioactive metabolites, and/or should depict antimicrobial effects for pathogens. Most importantly, it should be safe for further application (Navarré et al., 2024). Although many potential probiotic strains have been identified, their strain-specific behaviors, safety issues, and lack of reproducibility in animal studies hold their future application (Dash et al., 2023). Therefore, the search for novel probiotics is in constant demand. Moreover, studies on gut-inflammatory diseases have shown that the general physiological benefits of probiotics are strain-specific (McFarland et al., 2018). Therefore, the selection of novel probiotic strains requires well-rounded sequential evaluations for crucial probiotic properties.

Probiotics originating from the intestinal microbiota have a more preventive effect on host health than probiotics isolated from other sources (Timmerman et al., 2006). *Lactobacillus salivarius* is such a LAB that is majorly found in the oropharyngeal-gastrointestinal tract (OGT), milk, vagina, and oral cavities of humans (Yang et al., 2024; Guerrero Sanchez et al., 2022). *Lactobacillus salivarius* was recently reclassified from the *Lactobacillus* genus to a new genus called *Ligilactobacillus (*here onwards we will follow the same), which refers to its vertebrate host (Guerrero Sanchez et al., 2022). Being a native of the intestinal microbiota, *L. salivarius* helps to maintain a healthy and balanced gut-microbiome with multiple health-benefits (Yang et al., 2024). However, despite their promising potential, compared to other LAB, in-depth studies on *Ligilactobacillus salivarius* strains are scarce (Jiang et al., 2023) (**Supplementary Figure S1**). Here, we present well-rounded genomic, metabolomic, and functional studies of one such novel strain.

Whole genome sequencing (WGS) and phylogenetic analysis confirm the studied strain *LS-ARS2* as *Ligilactobacillus salivarius*. WGS analysis with the *LS-ARS2* genome predicts the presence of probiotic-specific gene signatures, which are functionally validated to establish the probiotic attributes (acid and bile tolerance, autoaggregation, cell-surface hydrophobicity, and cell-adhesion) of the strain. Interestingly, *LS-ARS2* appears as a fast-growing probiotic. Fast growth rate is not so common among LAB and could facilitate the strain for faster colonization in the gut. Moreover, *LS-ARS2* shows robust biofilm formation within 24 h; such efficacy is rare among *Ligilactobacillus salivarius* strains, indicating the possibility of the strain for efficient and longer sustainability in the gut. Further, we show significant antioxidant properties of the strain, indicating that *LS-ARS2* intake could reduce oxidative stress and prevent gut-dysbiosis. Moreover, *LS-ARS2* shows remarkable antimicrobial activities for Gram-positive and Gram-negative multidrug-resistant clinical strains. Fascinatingly, *LS-ARS2*-CFS shows significant inhibition for pathogenic biofilm. Therefore, it is not surprising to find the presence of potential antimicrobial peptide/bacteriocin encoding gene clusters in the *LS-ARS2-*genome. Finally, our genomic prediction, in corroboration with metabolomic studies (HRMS analysis), illustrates that the strain secretes diverse health-promoting metabolites. Therefore, this study systematically establishes *LS-ARS2 as* a promising probiotic with the potential application as a food-supplement and biotherapeutic to improve gut-health.

## 2. Materials and Methods

### 2.1 Bacterial Culture

Pure culture of *LS-ARS2* was procured from Microbial Type Culture Collection (MTCC), Institute of Microbial Technology (IMTECH), Chandigarh, India. The strain was maintained on MRS agar and MRS broth (HiMedia) in anaerobic conditions.

### 2.2 Reaffirmation of the Novel Strain Using Whole Genome Sequencing

#### 2.2.1 Isolation of DNA and preparation of library

*LS-ARS2* was grown in MRS broth anaerobically for 12 h at 37 °C. The genomic DNA was isolated using a DNA mini kit (QiaAmp DNA Mini Kit (Cat# 51306)) and quantified by Qubit Fluorometer 3 using the Qubit dsDNA High Sensitivity Assay Kit (Invitrogen, Cat# Q32854). Then it was further processed for library preparation with the 5300 Fragment Analyzer (3.1.0.12) using ProSize data analysis software 4.0.0.3. DNA libraries were subjected to pair-end sequencing using the NovaSeq6000 platform (MedGenome, Bangalore, India) with a read length of 151 bp. The generated sequence data was assessed for quality control and processed to generate FASTQ files. The 16S rRNA gene sequencing was performed, and the obtained consensus sequence was analyzed with the NCBI-BLAST tool in comparison with similar sequences present in the repository database. Neighbor-joining (NJ) phylogenetic tree was constructed using MEGA11 to identify the strain (Tamura et al., 2021).

#### 2.2.2 Determination of the strain identity by whole genome sequencing and genome assembly

The whole genome of the strain was sequenced [NovaSeq6000 (MedGenome, Bangalore, India)]. Initially, the reads were refined for contamination with human DNA. According to the alignment to the human genome (around 10.06%-27.65%), the reads were filtered for further alignment with a reference genome. The fastq file generated was checked for parameters like distribution of sequence quality score, base quality score, GC content in the reads, average base content per read, over-represented sequences, PCR amplification issue, and adapters. These parameters assisted to ensure the precision of the sequencing results. The sequences of fastq files were trimmed according to the quality report to only retain high-quality sequences for further analysis. fastq mcf (1.04.803) was used for adapter trimming, a process in which adapter sequences are removed from the 3’ end of the reads. Adapter trimming eliminates the chances of inference (due to adapters) in the alignment of the reads to a reference. For reference-guided assembly, the adapter trimmed reads were mapped to the suggested reference species *Ligilactobacillus salivarius strain* 609_LSAL to get the coverage and assembled fasta. The aligned consensus fasta files were generated [Samtools tools (version 1.2)]. Further, the depth statistics and coverage were generated using bedtools (version 2.0) and an in-house Perl script. The reference-aligned reads were used to predict variants using GATK, and the variants were annotated using the snpEff. The annotation of the primarily assembled genome was performed by applying the Prokaryotic Genome Annotation System (Prokka version 1.14.6) to predict genes, CDS, etc. (Seemann, 2014; Li and Durbin, 2009). The whole genome data can be retrieved from NCBI with the Bioproject Number PRJNA1024881.

### 2.3 Evaluation of Probiotic Attributes of *LS-ARS2*

#### 2.3.1 Tolerance to acidic pH

An acid tolerance assay was performed, followed by Abouloifa et. al. (Abouloifa et al., 2020) with modifications. *LS-ARS2* overnight culture was inoculated (1% v/v) in MRS broth adjusted to pH 4 and pH 3 with HCl (1N, Merck), followed by incubation for 0, 1, 3, and 5 h at 37 °C (Lab Companion, Korea). MRS broth (pH 6.5) was a control. After each time point, the culture was serially diluted in PBS (pH 7.4), 100 µL culture of appropriate dilution was plated on MRS agar, and incubated under anaerobic conditions at 37 °C for 24 h. Colonies were counted, and the viable cells, or the biomass (Log_10_ CFU/mL), were calculated. Survival rate (%) = [biomass in pH 4 or 3 / biomass in control pH 6.5] × 100. The experiment was performed in triplicate.

#### 2.3.2 Tolerance to bile salt

A bile tolerance assay was investigated according to Abouloifa et. al. (Abouloifa et al., 2020) with modifications. 1% of overnight culture was inoculated in MRS broth containing 0.3% and 1% (w/v) bile salts (HiMedia). MRS without bile salt was a control. After incubation for 0, 1, 3, and 5 h at 37 °C, each culture was serially diluted in PBS (pH 7.4), and 100 µL of the appropriate diluted culture was spread on MRS agar plate, thereafter incubated for 24 h at 37 °C under anaerobic conditions. After 24 h, the biomass (Log_10_ CFU/mL) was calculated by counting the colonies. Survival rate (%) = [biomass in bile salt 0.3% or 1% / biomass in control] × 100. The experiment was performed in triplicate.

#### 2.3.3 Determination of self-aggregation property of *LS-ARS2*

The aggregation property was evaluated according to Fonseca et. al. (Fonseca et al., 2021) and Campana et. al. (Campana et al., 2017) with modifications. 1% overnight culture of *LS-ARS2* was sub-cultured in MRS broth under anaerobic conditions till OD_600_ = 0.5-0.6. The culture was then harvested at 5000 rpm, washed using PBS (pH 7.4) twice, and dissolved in the same (PBS, pH 7.4). The absorbance of the culture was adjusted to approximately 10^8^ CFU/mL (OD_600_ 0.25-0.26 ± 0.1, A_0_) (UV-1800 UV Spectrophotometer, Shimadzu, Japan), vortexed for 10 sec, and incubated for 1, 2, 3, 4, 5, 6, 12, and 24 h at 37 °C. After each time interval, the absorbance of the upper suspension was measured at 600 nm (A_time_). The auto-aggregation percentage was determined by: Auto-aggregation (%) = [1 – (A_Time_/A_0_) × 100]. *L. acidophilus* DDS1, a widely used probiotic in clinical studies, was used as a positive control. The experiment was performed in triplicate.

#### 2.3.4 Evaluation of cell surface hydrophobicity of *LS-ARS2*

The surface hydrophobicity of *LS-ARS2* was evaluated using the Bacterial Attachment to Hydrocarbons (BATH) method according to Rosenberg et al. (Rosenberg et al., 1980) with modifications. The overnight culture was washed with phosphate urea magnesium sulphate (PUM) buffer and dissolved in 10 mL PUM to reach OD_600_ (A_0_) of 0.8-0.9. The adjusted cell suspension (4.8 mL) and n-hexadecane (Sigma)/xylene (Merck) (0.8 mL) were mixed, followed by incubation for 10 min. The cell suspension was vortexed for 2 min and kept at 37 °C for 2 h for the separation of phases. The absorbance of the lower aqueous phase was measured at 600 nm (A). Cell surface hydrophobicity (H%) was calculated as: Hydrophobicity (%) = [1 – A/A_0_) × 100]. Probiotic strain *L. acidophilus* DDS1 was used as a positive control. The experiment was performed in triplicate.

#### 2.3.5 Determination of adhesion property of *LS-ARS2* using human colon adenocarcinoma cells (Caco-2)

Adhesion ability was studied using Caco-2 cell lines (ATCC), maintained in complete DMEM (Himedia), supplemented with 10% heat-inactivated FBS (Gibco) in 5% CO_2_ (Galaxy 48R, New Brunswick, Germany). 2×10^4^ Caco-2 cells/well were seeded in six-well tissue culture plates (Dash et al., 2023; Ayala et al., 2019). After 80% confluent, the monolayer was washed using PBS (pH 7.4) twice, followed by incubation in serum-free DMEM overnight. Overnight bacteria at MOI (100:1) were co-cultured with Caco-2 cells and incubated for 2 h in a 5% CO_2_ atmosphere at 37 °C. Next, cells were washed using PBS (pH 7.4) five times, trypsinized, and gently aspirated. After serial dilution, 100 µL of appropriate dilution was plated on MRS agar followed by incubation for 24 h at 37 °C under anaerobic conditions. The attachment efficiency was determined by dividing the average number of bacteria (colonies after 24 h) attached per Caco-2 cell in each well (CFU/cell). The experiment was performed in triplicate.

#### 2.3.6 Elucidation of adhesion properties of *LS-ARS2* using phase-contrast confocal microscopy

To investigate the adhesion property, Caco-2 cells were seeded on poly-L lysine-coated (Sigma) coverslips (HiMedia) in 35 mm plates using complete DMEM and incubated up to 80% confluency (Dash et al., 2023). The cell monolayer was then washed using PBS (pH 7.4) followed by overnight incubation in serum-free DMEM. The bacteria at MOI (100:1) were co-cultured along with Caco-2 cells for 2 h, and unattached bacteria were washed by PBS (pH 7.4). The bacteria attached to Caco-2 cells were fixed with 4% paraformaldehyde for 30 min at 37 °C, followed by washing with water. 15 µL of mounting solution was placed on a glass slide, and the cover slip was placed on the mounting solution upside down. Then the cells were dried at room temperature, and finally, images were taken in a confocal microscope (FV3000, Olympus, Japan).

#### 2.3.7 Evaluation of biofilm-forming ability of *LS-ARS2*

The biofilm-forming ability of *LS-ARS2* was investigated as described earlier with minor changes (Rocchetti et al., 2024; Baldassarri et al., 2006). Briefly, the *LS-ARS2* strain was grown in MRS broth for 18 h under anaerobic conditions. Then the culture was diluted in MRS broth till OD_600_ = 0.1 and distributed in 24-well plates. The plates were incubated anaerobically for 24 h at 37 °C in a moist chamber. After the incubation period, the media was discarded, and the wells were washed with PBS (pH 7.4), and dried for 1 h at 60 °C. The remaining bacterial cells were stained for 45 min with 0.1% (w/v) crystal violet (SRL) solution (dissolved in 95% ethanol), washed with PBS (pH 7.4), and air-dried. The stain was dissolved in 33% acetic acid (Merck), and absorbance was measured at 570 nm. Probiotic strains *L. acidophilus* DDS1 and *L. rhamnosus* GG were used as positive controls. The assay was performed in triplicate.

### 2.4 Determination of Safety Attributes of *LS-ARS2*

#### 2.4.1 Prediction of prophage, CRISPR sequences, and antibiotic-resistant genes using genome analysis

PHAge Search Tool Enhanced Release (PHASTER) was implemented for the characterization of prophage sequences in the genome of *LS-ARS2* (Arndt et al., 2016). CRISPRCasFinder 1.1.2 (Grissa et al., 2007) was applied to screen CRISPR, Cas sequences, and truncated Cas sequences. The Comprehensive Antibiotic Resistance Database (CARD 3.3.0) (Jia et al., 2016) and the Resistance Gene Identifier Tool (RGI 6.0.3) were employed to find out the genes responsible for antibiotic resistance in the strain sequence using the criteria of perfect and strict hit and high-quality coverage (Zankari et al., 2012). The ResFinder 4.4.2 server was adopted to recognize the genes associated with acquired antimicrobial resistance with a threshold for %ID selected as 90.00% and minimum length selected as 60% and/or chromosomal mutations (Bortolaia et al., 2020).

#### 2.4.2 Evaluation of antibiotic susceptibility of *LS-ARS2*

Susceptibility of *LS-ARS2* to antibiotics was checked by the disc diffusion pattern following Fonseca et al. (Fonseca et al., 2021). Overnight culture was inoculated on MRS agar plates, and discs containing antibiotics (HiMedia) were placed. Inhibition zone diameters (mm) were measured after incubation for 24 h at 37 °C under anaerobic conditions. The susceptibility of the strain to the antibiotics was categorized as resistant (R), intermediate susceptible (I), or susceptible (S) according to CLSI guidelines (CLSI, 2020). The experiment was performed in triplicate.

#### 2.4.3 Assessment of haemolytic activity of *LS-ARS2*

Overnight culture was spot-inoculated on a sheep blood agar plate (defibrinated, 5% w/v, HiMedia) and incubated for 24 h at 37°C (Fonseca et al., 2021). The hydrolysis of blood cells resulted in a clear zone surrounding the colonies, which was considered β-hemolysis, whereas green-hued zones surrounding colonies designated partial hydrolysis, and were considered α-hemolysis, finally, no zone surrounding colonies was considered γ-hemolysis or non-haemolytic. *Staphylococcus aureus* ATCC 25923 was employed as a β-hemolysis positive control, *Escherichia coli* ATCC 25922 as a positive control for α-hemolysis, and reference probiotic strain *L. acidophilus* DDS1 for γ-hemolysis. The experiment was performed in triplicate.

#### 2.4.4 Estimation of DNase activity of *LS-ARS2*

Overnight culture was spot-inoculated on a DNase agar plate (HiMedia) and incubated at 37°C for 24 h (Fonseca et al., 2021). *Staphylococcus aureus* ATCC 25923 was considered a positive control, and the reference probiotic strain *L. acidophilus* DDS1 as a negative control for DNase activity. After incubation, the plates were flooded with 1 N HCl to develop a clear zone around the DNase producer colony. No clear zone around colonies was considered as negative DNase activity. The experiment was performed in triplicate.

#### 2.4.5 Evaluation of safety properties of *LS-ARS2* for human colon adenocarcinoma cells HCT116

The viability of *LS-ARS2*-treated colon cancer cells HCT116 (ATCC) was checked by the trypan blue exclusion assay based on Pradhan et al. with modifications (Pradhan et al., 2016). Briefly, 1×10^5^ HCT116 cells/well were seeded in complete DMEM (Himedia) supplemented with 10% heat-inactivated FBS (Gibco), and after attachment, the cells were treated with *LS-ARS2* at 100 MOI and incubated for 6, 12, and 24 h. The media was replenished every 3-4 h to avoid deprivation of the nutrients. After each time point, the cells were harvested and mixed with the same volume of 0.4% trypan blue (HiMedia). HCT116 without *LS-ARS2* treatment was considered a negative control. The living and the dead cells were counted. Cell viability was calculated as the number of viable cells divided by the total number of the cells. As a reference, probiotic strain *L. acidophilus* DDS1 was used. The experiment was performed in triplicate.

#### 2.4.6 Assessment of safety properties of *LS-ARS2*-derived CFS in the stationary phase of Caco-2 cells

Cytotoxicity of *LS-ARS2*-derived CFS was assessed on Caco-2 cells according to Pereira et al. with modifications (Pereira et al., 2022). *LS-ARS2* was grown in MRS broth followed by transfer in complete DMEM for 18 h in static condition at 37 °C. The culture was then centrifuged at 4000 rpm for 20 min at 4 °C, followed by filtration through a 0.22 µm filter to collect the CFS. Briefly, 2 × 10^4^ Caco-2 cells/well were seeded in 96-well plates in complete DMEM and allowed to grow for 7 days, with renewal of the media every 48 h. The confluent monolayer of Caco-2 in the stationary phase mimics the normal intestine cells. Hence, the viability of the cells after treatment assessed the safety of *LS-ARS2*-CFS for the host cells (Pereira et al., 2022). At day 7, cells were treated with different concentrations of CFS and pH-neutralized CFS (10%, 20%, 30%, 40%, 50%, and 60% CFS, diluted with complete medium) and incubated for 24 h. Caco-2 cells without treatment with CFS were considered a negative control. After the incubation period, 10 µL of MTT reagent (Sigma) was added and incubated for 4 h. Then, 100 µL DMSO (SRL) was added, and MTT absorbance was measured at 570 nm with background correction at 670 nm on the Spectramax iD3. The experiment was performed in duplicate with two technical repeats.

### 2.5 Evaluation of Antioxidant Properties of *LS-ARS2*

#### 2.5.1 Preparation of the intact and heat-lysed cells

Overnight cultures were washed by PBS (pH 7.4) twice. The cell pellet was re-suspended in distilled water to a final OD_600_ = 1.0. Heat-lysed cells were prepared by heating samples at 95 °C (water bath) for 30 min (Ramalho et al., 2019).

#### 2.5.2 Assessment of ABTS cation radical scavenging capacity of *LS-ARS2*

The ABTS cation radical scavenging assay was implemented as stated by Kim et al. (Kim et al., 2022) with minor changes. Working solution of ABTS (Sigma) was prepared by combining 7 mM ABTS and 2.45 mM potassium persulphate (HiMedia) in equal volume (1:1 v/v). The working solution was diluted with methanol (HiMedia, HPLC) up to OD_734_ = 0.7. 0.6 mL of samples (intact cells or heat-lysed cells) were mixed with 1.2 mL of ABTS solution and incubated for 30 min (dark, room temperature). Distilled water was used as a control. ABTS scavenging rate (%) = (Ac – As)/Ac × 100, where A_C_ is the absorbance of the control and A_S_ is the absorbance of the test sample measured at 734 nm. Probiotic strain *L. acidophilus* DDS1 was considered a positive control. The experiment was performed in triplicate.

#### 2.5.3 Determination of DPPH free radical scavenging capacity of *LS-ARS2*

The DPPH free radical scavenging assay was carried out following Ramalho et al. (Ramalho et al., 2019) with minor modifications. 1 mL of sample (intact cells or heat-lysed cells) was mixed with 1 mL 0.2 mM DPPH (CDH) solution in methanol (HiMedia). Mixed vigorously and incubated for 30 min (dark, room temperature). Distilled water was used as a control. DPPH radical scavenging activity (U/mL) = ABS_C_ – ABS_S_/S × 100, where ABS_C_ and ABS_S_ stand for the absorbance of the control and the test samples measured at 517 nm, respectively, and S is the sample volume (mL). Reference probiotic strain *L. acidophilus* DDS1 was considered a positive control. The experiment was performed in triplicate.

#### 2.5.4 Evaluation of superoxide anion scavenging activity of *LS-ARS2*

The superoxide anion scavenging activity was evaluated according to Gao et al. (Gao et al., 2013) with minor modifications. 0.8 mL of the sample (intact cells or heat-lysed cells) was added to 0.2 mL of Tris-HCl (Sigma) solution (0.1 M, pH 8). Subsequently, 0.1 mL of pyrogallol (3 mM, Sigma) was added, mixed, and incubated in the dark at room temperature (25 °C) for 30 min. The control group was taken as an equal volume of deionized water. Superoxide anion radical scavenging ability (%) = [1 – (A_s_ – A_1_)/A_0_] × 100, where A_s_ = the absorbance (320 nm) of the sample with pyrogallol, A_1_ = the absorbance of the sample solution lacking pyrogallol, and A_0_ = the absorbance of the blank solution with pyrogallol. Reference probiotic strain *L. acidophilus* DDS1 was considered a positive control. The experiment was performed in triplicate.

#### 2.5.5 Estimation of hydroxyl radical scavenging ability of *LS-ARS2*

The hydroxyl radical scavenging assay was evaluated according to Xiong et al. (Xiong et al., 2019) with minor modifications. Equal volumes of 2.5 mM 1,10-phenanthroline (Sigma), 0.2 M sodium phosphate buffer (Sigma), and FeSO_4_ (2.5 mM, Merck) were vortexed and incubated for 5-7 min. Then an equal volume of H_2_O_2_ (0.12% v/v, CDH) and sample (intact cells or heat-lysed cells) were added, vortexed, and incubated in a water bath at 37 °C for 60 min. The control group was taken as an equal volume of deionized water instead of the sample. Hydroxyl radical scavenging activity (%) = [(A_s_ – A_1_)/(A_0_ – A_1_)] × 100, where A_s_ = the absorbance (536 nm) of the sample containing H_2_O_2_, A_1_ = the absorbance of the sample deprived of H_2_O_2_, and A_0_ = the absorbance of the solution with H_2_O_2_ and without a sample. Reference probiotic strain *L. acidophilus* DDS1 was considered a positive control. The experiment was performed in triplicate.

### 2.6 Determination of antimicrobial properties of *LS-ARS2*

#### 2.6.1 Preparation of the cell-free supernatant (CFS)

*LS-ARS2* was grown anaerobically in MRS broth at 37 °C for 24 h. The culture was centrifuged at 4000 rpm for 20 min at 4 °C. The supernatant was sterilized by filtration through a 0.22 µm PVDF filter (HiMedia) and used for the experiments (Chen et al., 2019).

#### 2.6.2 Evaluation of the antimicrobial activity of *LS-ARS2* using Gram-positive and Gram-negative pathogens, including muti-drug resistant hospital strains

The antimicrobial effect was assessed for enteric pathogens by the agar well diffusion assay according to Abouloifa et al. (Abouloifa et al., 2020) with modifications. The studied pathogens include enteric Gram-negative strain *Salmonella typhimurium* ATCC 14028, Gram-positive Methicillin-resistant *Staphylococcus aureus* ATCC 700699 (MRSA)*, Staphylococcus aureus* ATCC 25923, as well as clinically isolated Gram-negative multi-drug-resistant strains like *Escherichia coli* (ETEC) BCH 04067*, Vibrio cholerae* BCH 09616, and *Shigella Flexneri* BCH 06745. The pathogenic strains were swabbed at a concentration of 0.5 McFarland (OD_600_ = 0.5) on Mueller Hinton agar (HiMedia), and 6 mm wells were made on agar medium. The wells were loaded with 100 μL CFS of *LS-ARS2*. The antimicrobial activity was determined by measuring the clear zone of inhibition around the wells in MHA plates after overnight incubation at 37 °C. For the negative control, MRS broth (uninoculated) was used. *L. rhamnosus* GG (*LR-GG*), an established probiotic, was used as a positive control. The experiment was performed in triplicate.

#### 2.6.3 Co-culture experiments of *LS-ARS2* with *Salmonella typhimurium*

The antibacterial effect of *LS-ARS2* was further validated by co-culture assay with *Salmonella typhimurium* (ATCC 14028) according to Chen et al. (Dash et al., 2023) with slight modifications. Overnight cultures of *LS-ARS2* and *S. typhimurium* were centrifuged and washed with PBS (pH 7.4). First, different doses of *LS-ARS2* (10^6^, 10^7^, and 10^8^ CFU/mL, respectively) were co-cultured with the pathogen (10^6^ CFU/mL) till 24 h, and the viable cells were counted on SS agar plates (HiMedia). To perform the time-kill assay, *S. typhimurium* was co-cultured together with *LS-ARS2* in a 1:1 ratio (10^6^: 10^6^ CFU/mL) for multiple time points (0, 3, 6, 12, 24 h), and respective colonies of *S. typhimurium* (ST) were counted. ST without *LS-ARS2* was considered a control. The experiment was performed in triplicate.

#### 2.6.4 Calculation of minimum inhibitory percentage (MIP) of *LS-ARS2* CFS using broth microdilution method

For the determination of MIP, a microdilution assay was implemented according to Chen et al. (Chen et al., 2019). The overnight culture of *S. typhimurium* was centrifuged and washed by PBS (pH 7.4) twice. CFS was used in three sets: set 1: used as it is; set 2: heated at 95 °C for 15 min; set 3: pH of the CFS was neutralized using NaOH. 100 µL of Mueller Hinton broth (MHB, HiMedia) comprising 10^5^ CFU/mL *S. typhimurium* was mixed with different percentages (i.e., 1%, 5%, 10%, 20%, 30%, 40%, 50%) of the CFS diluted in MRS in a microtiter plate. Un-inoculated MHB and MRS were applied as negative controls (blank). The plates were incubated for more than 18 h, and the absorbance was measured at 600 nm using a microplate reader. From the same microtiter plate, cultures were spot-inoculated on MHA plates to evaluate bacteriostatic or bactericidal effects on the pathogen. The standard probiotic *LR-GG* was used as a positive control. The experiment was performed in triplicate.

#### 2.6.5 Evaluation of anti-biofilm properties of *LS-ARS2* CFS

The antibiofilm property of cell-free supernatant of *LS-ARS2* was assessed by the crystal violet staining method according to Lee et al. with modifications (Lee et ^a^_l., 202_^1^). Overnight-grown MRSA strain was adjusted to OD_600_ = 0.1 in Tryptone Soya Broth (TSB, HiMedia) and dispensed into each well of 24-well plates with 100 mM glucose (Merck) and NaCl (Merck). CFS of *LS-ARS2* was added to the pathogen at a range of concentrations of 1.25%-40% v/v (1/8 MIP to 4 MIP). After incubation at 37 °C for 24 h, the medium was removed, and staining was performed as mentioned above (section 2.3.7.). Un-inoculated MRS with TSB broth was used as a control. The biofilm formation rate (%) was calculated according to the following formula: Biofilm formation rate (%) = (OD_Sample_/ OD_Control_) × 100. The experiment was performed in triplicate.

#### 2.6.6 Prediction of bacteriocins using the whole genome sequence of *LS-ARS2*

Biosynthetic gene clusters (BGCs) are closely associated groups of genes on a genome that together work to synthesize specialized metabolites. The BGCs of antimicrobial compounds (bacteriocins) of the *LS-ARS2* genome were explored by the web server, BAGEL4 (de Jong et al., 2006).

### 2.7 Determination of functional annotations of the *LS-ARS2* genome

The prediction of genes in the *LS-ARS2* genome was performed using Prokka (1.14.6), and the functional annotation was done using Emapper. The cluster of orthologous groups (COG) intended for the protein-coding genes was determined by Egg-NOG mapper (version 2.1.12) (Dietzsch et al., 2021) using the online Egg-NOG database (version 5.0). The presence of carbohydrate-active enzyme (CAZymes) encoding genes in the *LS-ARS2* genome was explored with the meta server, dbCAN3 (Zheng et al., 2023).

### 2.8 Prediction of primary and secondary metabolites using the whole genome sequence of *LS-ARS2*

The *LS-ARS2* genome sequence was analyzed in gutSMASH (Specialized Primary Metabolite Analysis from Anaerobic Bacteria) for the prediction of potential primary metabolites (Pascal Andreu et al., 2021). Similarly, the secondary metabolites of the strain were predicted using antiSMASH version 6.0.1 (Antibiotic and Secondary Metabolites Shell) using strictness “strict” (Blin et al., 2021).

### 2.9 Elucidation of metabolite profiling of *LS-ARS2* by High-Resolution Mass Spectroscopy (HRMS)

The metabolite composition of the *LS-ARS2*-derived CFS was analyzed using an Exactive™ Plus Orbitrap high-resolution mass spectrometer linked in tandem to an Ultimate 3000 high-performance liquid chromatography column (Thermo Scientific, USA) (Dash et al., 2023). Liquid chromatography-mass spectrometry (LC–MS)-grade water, methanol, acetonitrile, and formic acid were used (Fisher Scientific, USA). Equal volumes of CFS and methanol (1:1 ratio) were methodically combined and sterilized by syringe filtration using a 0.22 μm filter. For mass spectrometry, around 0.5 mL of the filtered solution was added to the DP ID vial (Cat# C4000-1W, Thermo Scientific, USA). The components in the sample were sorted out by a C18 column (Hypersil BDS, 250 mm × 2.1 mm, 5 μm; ThermoScientific, USA). The mobile phase contained acetonitrile and water (1:1) containing formic acid (0.1%). The flow rate of the sample was adjusted to 3 μL/min, the temperature of the column was set at 30 °C, and the pressure was 700 bar. Electrospray ionization at a potential of 3 eV was used to ionize the molecules. The sample run time was 5 min, and both ions (positive and negative) were detected within the 50-750 m/z scan range. Metabolite annotation of the mass peaks was performed through the National Metabolomics Data Repository (NMDR). The peaks were then annotated with the Metabolomics Workbench database (https://www.metabolomicsworkbench.org/data/M_form.php#M2) using public sources such as LIPID MAPS, ChEBI, HMDB, BMRB, PubChem, NP Atlas, and KEGG. For compound consolidation, mass tolerance (the acceptable deviation from the obtained m/z) was set at 0.2 ppm. The results of the analysis were exported with the following filtration criteria: delta PPM ranges from -0.2 ppm to +0.2 ppm and the complete annotated name of the compound. Both positive and negative mode analyses were performed simultaneously, which were combined at a later stage, and duplicates were removed. The analyzed molecules were clarified on the basis of the published literature and recorded in the table.

### 2.10 Statistical analysis

Statistical analysis was performed by two-way ANOVA (for comparison between more than two groups and two parameters) and one-way ANOVA (for comparison between more than two groups and one parameter) to determine significant differences between the treatments using GraphPad Prism (Version 8). The differences were considered statistically significant between all the treatment groups when p≤0.05.

## 3. Results

### 3.1 Whole genome sequencing features of *LS-ARS2*

Our whole genome sequence study revealed that the genome of *LS-ARS2* was a single circular chromosome. The complete genome sequence contained 1,810,531 bp. Average GC content was found to be 34.4% (**Figure 1A**). The coding DNA sequences (CDSs) were 1586, and the identified genes were 1610 (**Table 1**). The chromosome contained 4 rRNAs, 19 tRNAs, and 1 tmRNA encoding sequence. The overall summary of the genome assembly was included in **Supplementary Table S1**.

**FIGURE 1.**
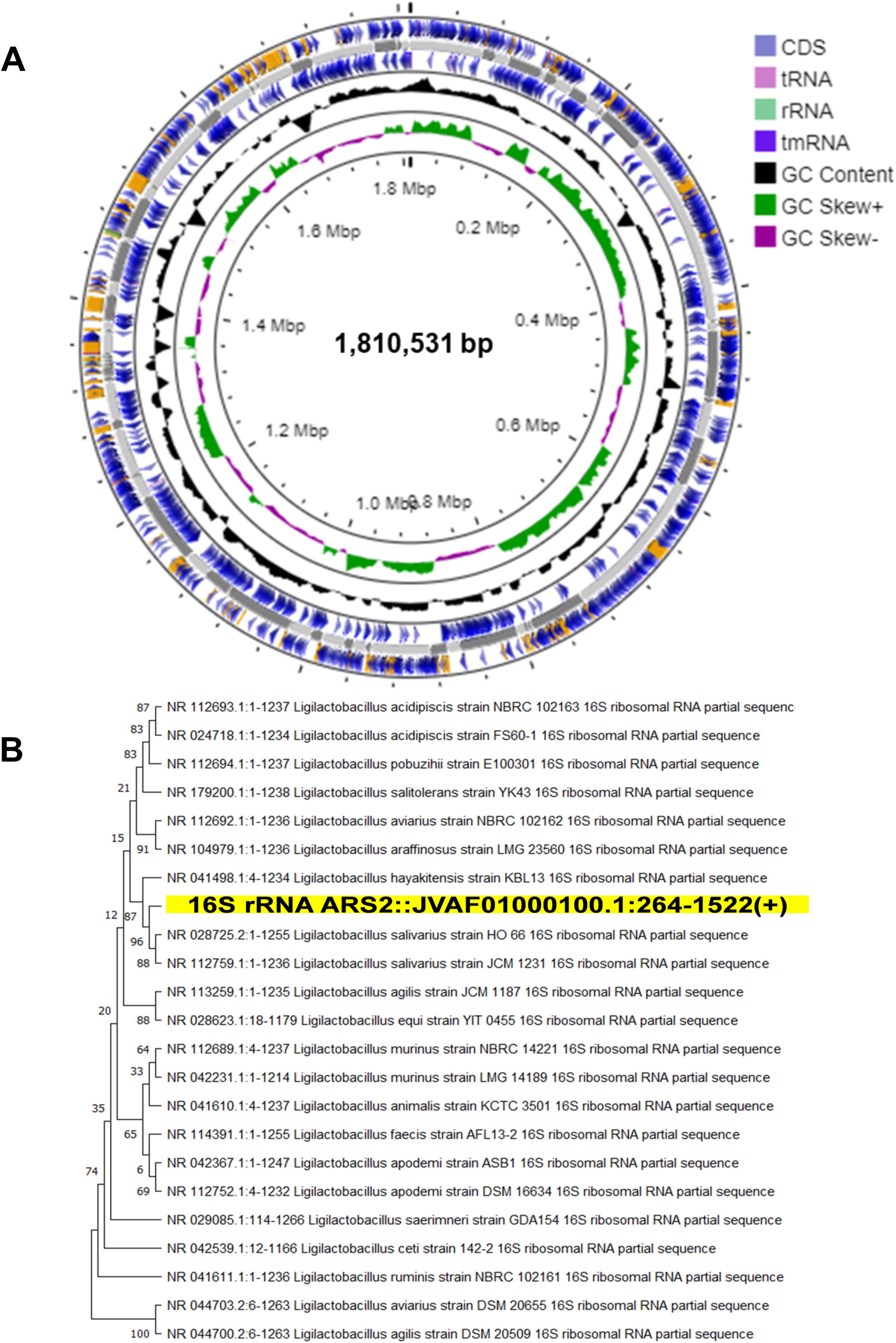
(A) Circular genome diagram of *LS-ARS2*. From the inner to the outer: the first ring represented the genome size (1,810,531 bp); the second depicted the GC skew (G + C/G – C); the third and fourth showed forward and reverse CDS (coding sequences) annotated with Prokka (the sites of CDSs/rRNA/tRNA/tmRNA on the genome are marked). **(B) Phylogenetic analysis based on NCBI blast and MEGA 11 reaffirmed that *LS-ARS2* belongs to *Lactobacillus salivarius.*** Phylogenetic tree indicates that the closest clad for *LS-ARS2* is *Ligilactobacillus salivarius* strain HO 66.

**Table 1.** Complete genomic feature of *LS-ARS2*.

| **Strain** | ***LS-ARS2*** |
| --- | --- |
| Genome length (bp) | 1810531 |
| GC content (%) | 34.42 |
| Total number of genes | 1610 |
| Coding genes | 1586 |
| tRNA number of the assembled genome | 19 |
| rRNA number of the assembled genome | 4 |
| tmRNA number of the assembled genome | 1 |
| CRISPR-Cas array | 3 CRISPR and 5 Cas-associated arrays |
| Prophage (intact region) | 0 |
| Antibiotic acquired genes | NONE |

### 3.2 Phylogenetic analysis and taxonomic position reaffirmed the strain as *Ligilactobacillus salivarius*

A BLAST search was performed with all the *Ligilactobacillus* complete genomes to identify the species closest to our strain. Phylogenetic analysis showed the close relations of our strain with other species of the genus *Ligilactobacillus* and grouped in a single clade with *Ligilactobacillus salivarius* strain HO66 (**Figure 1B**). This phylogenetic analysis result corroborated with the average nucleotide identity (ANI) analysis, where 98% of ANI identity was found for *LS-ARS2* with *L. salivarius* DSM 2055 and 87.49% alignment coverage with *Ligilactobacillus salivarius* strain 609_LSAL. (Tenea and Ortega, 2021). Moreover, as per 16S rRNA extracted using Barrnapp, *Ligilactobacillus salivarius* strain HO 66 showed the highest identity with our strain. Based on this, our strain was inferred as *Ligilactobacillus salivarius*.

### 3.3 Probiotic Attributes of *LS-ARS2:* Genomic analysis coupled with *in vitro* investigations

#### 3.3.1 Survival potential of *LS-ARS2*: acid and bile tolerance

Probiotics are expected to survive the acidic environment of the GI tract to reach the small intestine, where they show beneficial effects. In addition to an acidic environment, probiotics must withstand high bile salts in the small intestine.

Our WGS analysis indicated the presence of several ABC transporters, sodium proton antiporter, sodium bile acid symporter family, ABC-type Na+ efflux pump, and permease component in the *LS-ARS2* genome **(Supplementary Table S2)**. Moreover, genes encoding universal stress protein, multi-chaperon systems (recover cells from stress-induced conditions), grpE (responds to hyperosmotic conditions), and heat shock proteins were identified in the *LS-ARS2* genome **(Figure 2A and Supplementary Table S2)**. Therefore, WGS analysis indicated that the strain harbors different stress-tolerance genes and proteins that might enable them for stress adaptation.

**FIGURE 2.**
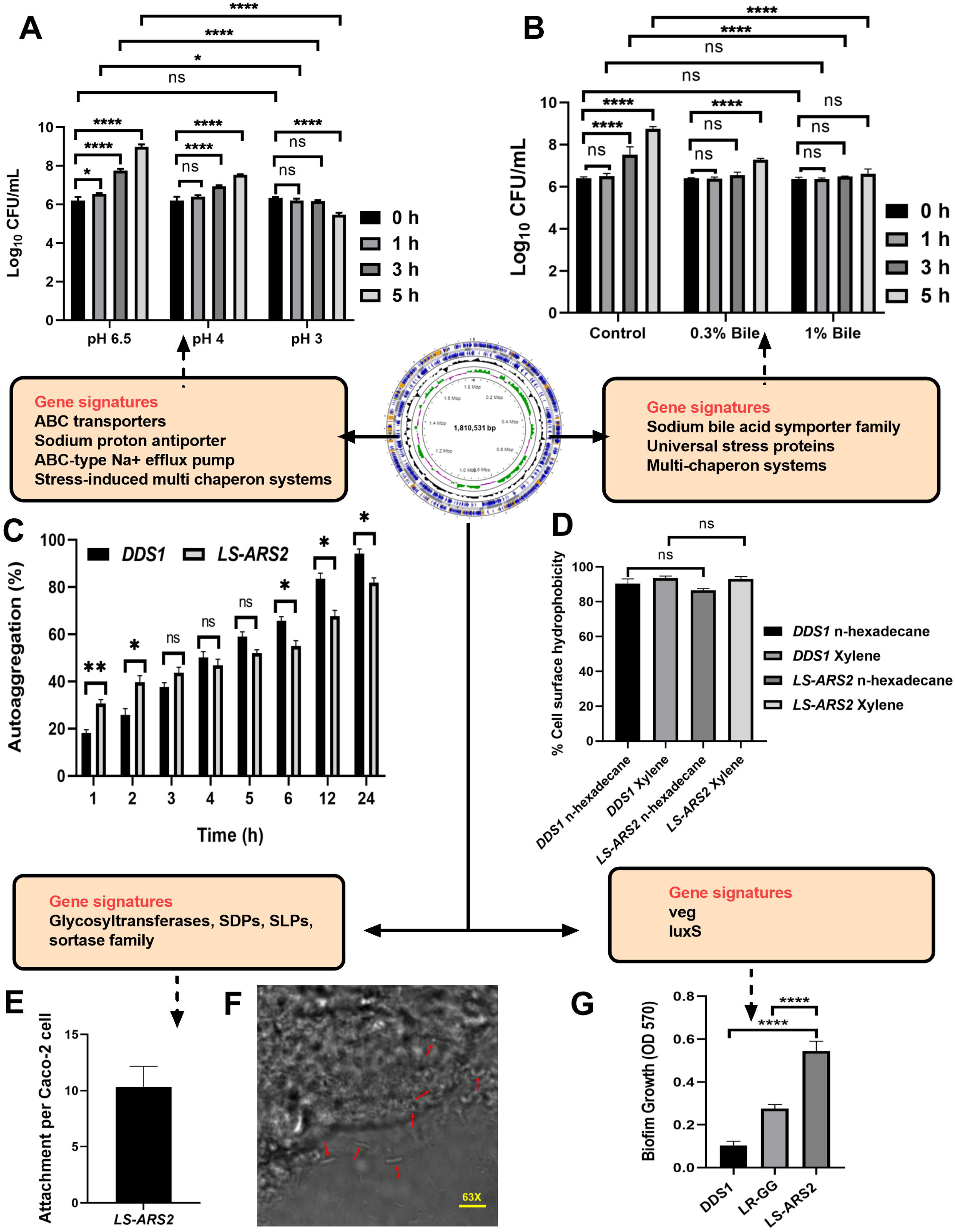
Probiotic properties of *LS-ARS2*. **(A) Acid tolerance of *LS-ARS2***. Survival of *LS-ARS2* at pH 3 and pH 4 at different time intervals indicated the viability of the strain at acidic pH. Data was presented as mean ± SD from three independent biological replicates. Statistical analysis was done using a two-way ANOVA with a Bonferroni post hoc test; *p<0.05; ****p<0.0001. **WGS analysis** indicated the presence of potentially related stress-tolerance genes like ABC transporters, sodium proton antiporter, ABC-type Na+ efflux pump, and permease in the *LS-ARS2* genome. **(B) Bile tolerance of *LS-ARS2.*** Viability of *LS-ARS2* in the presence of 0.3% and 1% bile salt for 3 and 5 h of incubation time indicated tolerance of the strain at high bile concentrations. Data was represented as mean ± SD based on three independent biological replicates. Statistical analysis was done using a two-way ANOVA with a Bonferroni post hoc test; *p<0.05; ****p<0.0001. **Genomic analysis** illustrated the presence of potentially related stress-tolerance genes like sodium bile acid symporter family genes, genes for universal stress protein, and multi-chaperon systems in the *LS-ARS2* genome. **(C) Autoaggregation attributes of *LS-ARS2.*** The aggregation property of *LS-ARS2* was assessed side by side and compared with the reference probiotic strain *L. acidophilus* DDS1 at different time points up to 24 h. The results were depicted as mean ± SD based on three independent biological replicates. Statistical analysis was done using two-way ANOVA with Bonferroni post hoc test; p>0.05. **(D) Cell surface hydrophobicity property of *LS-ARS2.*** The hydrophobicity percentage of *LS-ARS2 was* evaluated with n-hexadecane and xylene (2 h of incubation) in comparison to the reference strain *L. acidophilus* DDS1. All the results were shown as mean ± SD based on three independent biological replicates. Statistical analysis was done using one-way ANOVA with Bonferroni post hoc test; *p<0.05; ****p<0.0001. **(E) Adhesion efficacy of *LS-ARS2***. Attachment efficiency of *LS-ARS2* was shown with intestinal epithelial Caco-2 cells. **WGS analysis** (functional annotation) depicted the abundance of related genes like glycosyltransferases, genes for sortase-dependent proteins (SDPs), and S-layer proteins (SLPs) in the *LS-ARS2* genome **(F) Visualization of Adhesion property of *LS-ARS2.*** Phase contrast confocal microscopy images of the Caco-2 cells after co-culturing with *LS-ARS2* showed efficient adhesion to Caco-2 cells. **(G) Biofilm-forming ability of *LS-ARS2***. The assay was performed compared with two standard probiotic strains, *L. acidophilus* DDS1 and *L. rhamnosus* GG (*LR-GG*). Results were represented as mean ± SD based on three independent biological replicates. Statistical analysis was done using one-way ANOVA with Bonferroni post hoc test; ****p<0.0001. **WGS analysis** revealed that the *LS-ARS2* genome consists of *veg* and *luxS* genes, reported to be associated with biofilm formation in bacteria.

Further, acid tolerance assay showed *LS-ARS2* exhibited 80% and 61% survival in pH 3 after 3 and 5 h of incubation, respectively (**Figure 2A**). Interestingly, there was no significant difference in the biomass with control MRS up to 1 h of incubation for both pH conditions (p>0.05). Moreover, the viable count increased significantly (****p<0.0001) after 3 and 5 h of incubation at pH 4. However, in pH 3, the strain survived well up to 3 h (p>0.05), but the viable cells decreased significantly after 5 h of incubation (****p<0.0001). Therefore, *LS-ARS2* was able to survive in the acidic pH 3 and pH 4.

Next, we performed bile tolerance assay. The bile salt concentration varies from 0.2-0.3% depending on the type and amount of food ingested (Hsiung et al., 2021). The strain exhibited 87% and 83% survival in 0.3% (w/v) bile salt after 3 and 5 h of incubation, respectively. The strain also showed a survival of 86% and 76% in 1% (w/v) bile salt after the above-mentioned incubation time (**Figure 2B**). Interestingly, the survivability of the strain was similar to the control versus 0.3% or 1% bile salt up to 1 h (p>0.05). Moreover, no significant changes were observed in viable count between 0 and 3 h for both 0.3% and 1% bile salt (p>0.05). In addition, the biomass was found to increase significantly (****p<0.0001) after 5 h in the presence of 0.3% bile. Therefore, the strain could potentially be able to survive at high bile salt.

In summary, our genomic analysis together with the results from biochemical assays strongly supported the survival of the strain *LS-ARS2* in harsh environments in the GI tract.

#### 3.3.2 Colonization potential of *LS-ARS2:* Self-aggregation, cell surface hydrophobicity property, and adhesion efficacy in Caco-2 cells

Probiotics are expected to colonize in the gut and form a strong barrier on the epithelial mucosa that restricts the entry of pathogens.

Functional annotation of the genes by COG analysis exhibited the abundant presence of glycosyltransferases in the *LS-ARS2* genome that might provide a specific benefit to adhere and colonize in the GI tract (Dash et al., 2023). The envelope of *Lactobacilli* contains many cell-surface proteins involved in *lactobacillus*-host interactions. Among them, sortase-dependent proteins (SDPs) and S-layer proteins (SLPs) are best characterized for providing adhesion properties (Lebeer et al., 2008). In the present study, srtA (a sortase family protein) and lspA (involved in adhesion through prolipoproteins) were found by genome COG analysis supporting the adhesion potential of *LS-ARS2*. The genome of *LS-ARS2* also contained several cell wall biosynthetic proteins such as dltD, dltC, dltA, dltB, and dltX, which might influence adhesion capacity and immune response in the host cells (Dash et al., 2023; Lebeer et al., 2008).

Since aggregation of microbes is important for gut-adhesion and colonization, evaluation of self-aggregation is considered an important selection criterion for potential probiotics. Our strain showed a significant increase in autoaggregation capacity from 30.6% to 67.6% within 1-12 h, while it showed the highest autoaggregation ability (81.75%) after 24 h (**Figure 2C**). The auto-aggregation ability shown by the reference probiotic strain *L. acidophilus* DDS1 was also comparable to *LS-ARS2* (^ns^p>0.05, *p<0.05, **p<0.05).

Surface hydrophobicity of the bacterial cells is also pivotal for gut-colonization (Hsiung et al., 2021). *LS-ARS2* showed significant surface hydrophobicity with n-hexadecane (86.59%) and xylene (93%) (**Figure 2D**). The hydrophobicity of *LS-ARS2* was similar to the reference strain *L. acidophilus* DDS1 (p>0.05). This result indicated the potential adhesion capacity of the strain with complex hydrophobic substratum.

One of the most important properties of probiotics is their capacity to adhere to the gut epithelial cells. Therefore, next, we wanted to study directly the adhesion potential of the strain for colon epithelial cells (Caco-2). Significant adhesion capacity (10-12 of *LS-ARS2*/ well-differentiated Caco-2 cells) was observed after 2 h of incubation (**Figure 2E**). The representative phase-contrast confocal microscopy images also reaffirmed the substantial attachment of *LS-ARS2* to Caco-2 cells. Therefore, the strain indeed showed promising adhesion ability for the human gut lining (**Figure 2F)**.

Therefore, genomic analysis together with *in vitro* assays indicate the potential of the strain *LS-ARS2* to colonize in the gut.

#### 3.3.3 Biofilm forming ability of *LS-ARS2*

Biofilm formation is reported to be a beneficial property of *Lactobacillus* sp. since the formation of biofilm showed better colonization, survival, and prolonged persistence in the gastrointestinal mucosa in the host (Salas-Jara et al., 2016). Probiotic-derived biofilm was also reported to secrete exopolysaccharides, vitamins, enzymes, and many metabolites to enhance the growth of gut-microbiota and inhibit pathogenic adhesion (Gao et al., 2022).

Biofilm-forming ability among *L. salivarius* strains is rare and strain-specific. However, our WGS analysis of *LS-ARS2* revealed the presence of *veg* and *luxS* genes in the *LS-ARS2* genome. It is worth mentioning that *veg* is a widely reported stimulator for biofilm formation in Gram-positive bacteria (Lei et al., 2013). Further, through quorum sensing, *luxS*-derived autoinducer-2 (AI-2) plays a crucial role in adhesion and formation of biofilm in *Lactobacillus spp*. [Mgomi et al. 2023; Liu et al. 2018]. Therefore, the presence of the *veg* and *luxS* genes in the *LS-ARS2* genome encouraged us to explore the biofilm-forming ability of *LS-ARS2* experimentally.

It was extremely exciting to find that *LS-ARS2* was able to form robust biofilm within 24 h without any additional carbon source or supplements. Moreover, biofilm formation by *LS-ARS2* was found to be more robust than the reference probiotic strains DDS1 and *LR-GG* (****p<0.0001) (**Figure 2G**). *LS-ARS2*-biofilm could therefore offer a promising application potential of the strain for improving gut-health.

### 3.4 Safety attributes of *LS-ARS2*: WGS analysis and *in vitro* validation

#### 3.4.1 Genome stability of *LS-ARS2*

Numerous *Lactobacillus* species contain hidden prophages with the potential of cross-contamination or modulation of the intestinal microecology when released (Pei et al., 2021). Therefore, it is essential to perform a comprehensive scanning of prophages in probiotic candidates for their safety evaluation. While analyzing the genome sequence of *LS-ARS2*, we found four prophage regions (PHASTER). However, all of them were incomplete prophage regions (**Supplementary Table S3**). Moreover, bacteria harbour adaptive antiphage defense mechanisms like the CRISPR/Cas system for their protection from phage attacks (Pei et al., 2021). In our strain, three CRISPR arrays and five Cas-associated sequences (CRISPRFinder) were predicted (**Supplementary Table S4**). Next, a previous study indicated that *Lactobacillus* genomes carrying type I or type III CRISPR-Cas systems harbored fewer intact prophages (Pei et al., 2021). Indeed, TypeIE Cas-associated systems were the most prevalent in the genome of our strain as well. Hence, the presence of a TypeI Cas-associated system in *LS-ARS2* might act as an antiphage defense system and could cause the absence of intact prophage regions.

Microorganisms may acquire antibiotic-resistant genes by horizontal gene transfer (Jia et al., 2016). Several *Lactobacillus* strains carry d-Ala-d-lactate in their peptidoglycan rather than the d-Ala-d-Ala dipeptide, which makes them resistant to vancomycin (Rychen et al., 2018). For example, *L. plantarum* strains were found with intrinsic vancomycin resistance genes (Deghorain et al., 2007). The genome of *LS-*

*ARS2* was also predicted to have one strict antibiotic-resistant gene (CARD database and RGI analysis, **Supplementary Table S5**). However, the same was not found with Resfinder analysis.

#### 3.4.2 Antibiotic susceptibility of *LS-ARS2*

Extensive use of antibiotics with food supplements can accumulate genes responsible for antibiotic-resistance in gut-bacteria. These antibiotic-resistant genes can be acquired by the pathogens sharing the corresponding intestinal niche, which appears as a clinical threat (Wong et al., 2015). Therefore, to project any probiotic as a food supplement, the assessment of the antibiotic susceptibility profile is crucial to ensure the safe application. In our study, *LS-ARS2* was sensitive to penicillin, erythromycin, clindamycin, tetracycline, ampicillin, chloramphenicol, and amoxicillin/clavulanic acid. However, it was found to be resistant to vancomycin and gentamicin, similar to many probiotic strains (**Table 2**). Therefore, *LS-ARS2* was sensitive to the majority of the conventional antibiotics.

**Table 2.** Antibiotic susceptibility of *LS-ARS2* strain determined by the disk diffusion method.

| Mode of action | Antibiotics | Results<br>*LS-ARS2* |
| --- | --- | --- |
| Protein Synthesis Inhibitors | Erythromycin (15 µg) | S* |
|  | Gentamicin (10 µg) | R*** |
|  | Chloramphenicol (30 µg) | S |
|  | Tetracycline (30 µg) | S |
|  | Clindamycin (2 µg) | S |
| Cell wall synthesis inhibitors | Penicillin (10 µg) | S |
|  | Amoxicillin/Clavulanic acid (20/10 µg) | S |
|  | Vancomycin (30 µg) | R |
|  | Cephalothin (30 µg) | S |
|  | Ampicillin (10 µg) | S |

✓ **Zone diameter (mm) Interpretive Criteria: $\geq 20$ = S\* (Susceptible); 15-19 = I\*\* (Intermediate); $\leq 14$ = R\*\*\* (Resistant) [CLSI guideline] [24]**
✓ **The result is presented from three individual biological replicates**

#### 3.4.3 Non-haemolytic and DNase negative properties of *LS-ARS2*

The non-haemolytic nature of *LS-ARS2* was signified by the nonappearance of any transparent zone at the blood agar plate. Further, our strain did not show any halo zone around the colonies at the DNase agar plate, which suggested the absence of DNase activity (**Table 3**). Therefore, *in silico* studies and *in vitro* assays assured the non-pathogenic nature of *LS-ARS2*.

**Table 3.** Haemolytic and DNase activity of *LS-ARS2*.

| Strains | Haemolytic Activity |  |  | DNase Activity |
| --- | --- | --- | --- | --- |
| | $\alpha$ | $\beta$ | $\gamma$ | |
| *E. coli* ATCC 25922 | + | - | - | NA |
| *S. aureus* ATCC 25923 | - | + | - | + |
| *L. acidophilus* DDS1 | - | - | + | - |
| *LS-ARS2* | - | - | + | - |

✓ *L. acidophilus* DDS1, a widely used probiotic in clinical studies, is considered as a reference strain.
✓ The result is presented from three individual biological replicates

#### 3.4.4 Safety attributes of *LS-ARS2* in human cell lines

To further secure the safety attributes of *LS-ARS2*, both the whole cell and cell-free supernatant of the strain were tested on human colon cancer cells. The trypan blue exclusion assay ensured that the application of *LS-ARS2* does not show any notable toxicity to HCT116. A similar result was also observed for the standard probiotic strain *L. acidophilus* DDS1 (**Figure 3A and 3B**). In the MTT assay, the CFS of *LS-ARS2* also did not show any notable toxicity as compared to the control (complete DMEM without CFS) to the stationary-phase Caco-2 cells, which mimic the normal intestinal cells (**Figure 3C and 3D**). MTT and trypan blue exclusion assays therefore ensured the safe application of the strain on human cells.

**FIGURE 3.**
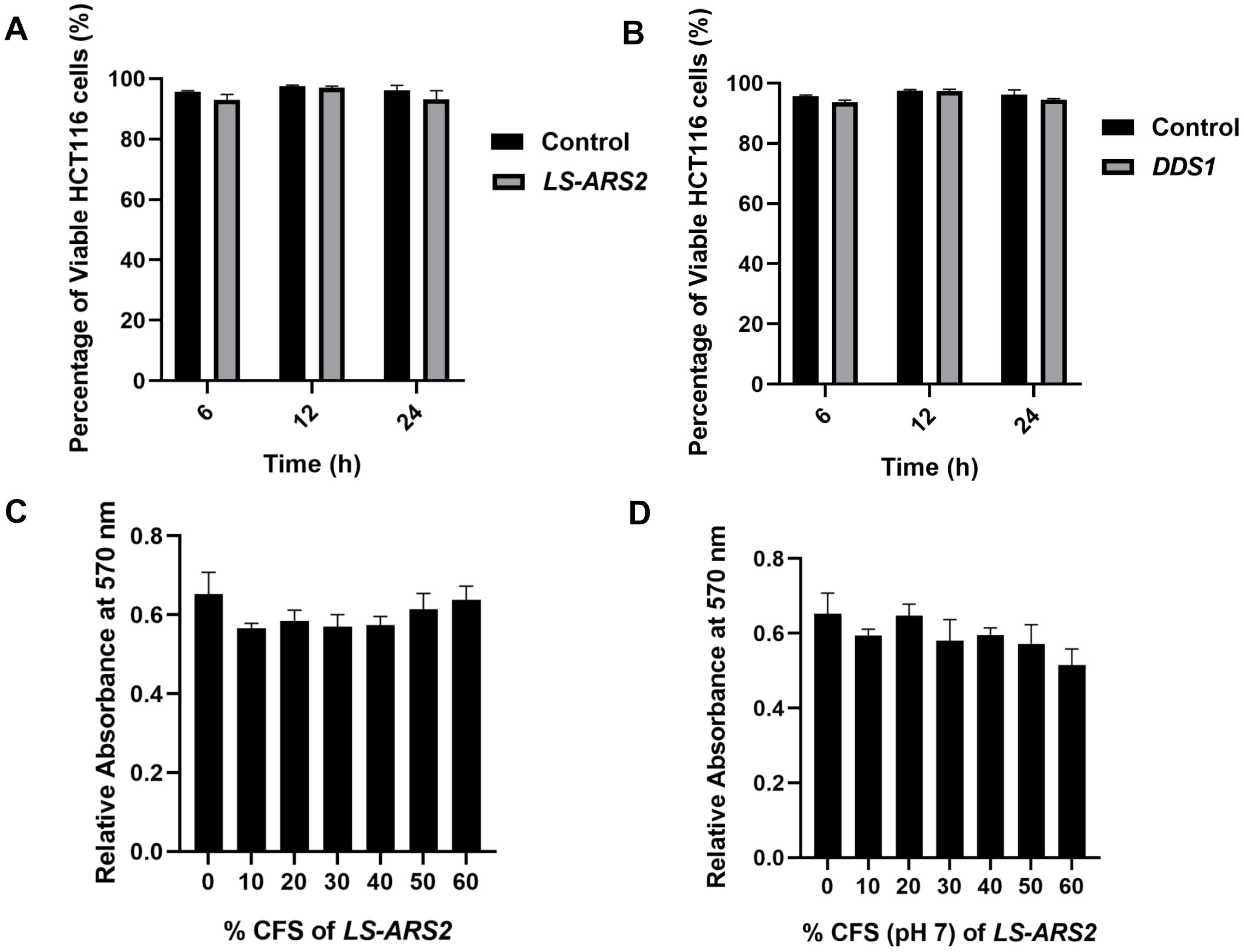
(A-B) Trypan blue exclusion assay. The result showed the percentage viability of human colorectal cancer cell HCT116 after treatment with **(A)** *LS-ARS2* and **(B)** *L. acidophilus* DDS1. All the data were represented as the mean ± SD based on three independent biological replicates. **(C-D) MTT assay. (C)** *LS-ARS2* CFS and **(D)** pH-neutralized CFS (pH 7) of *LS-ARS2* on Caco-2 cells showed non-toxic nature of *LS-ARS2-*derived CFS.

### 3.5 Functional attributes of *LS-ARS2:* Antioxidant potential of *LS-ARS2*

Probiotics with antioxidant potential could alleviate the oxidative stress in the host, hence could serve as a natural antioxidant. The functional annotation of the *LS-ARS2* genome annotated multiple genes responsible for antioxidant activities (**Supplementary Table S2**). The genes encoding whole thioredoxin (*trxa, trxb, tpx*) and NADH (*ndh, npr*) antioxidant systems were detected, which are reported to be associated with scavenging of reactive oxygen species (ROS), detoxifying peroxidases, and protecting cells from oxidative stress (Kandasamy et al., 2024; Zhang et al., 2018). Additionally, the genes encoding for glutathione synthetase (*gshF*) and glutaredoxin (*nrdH*) were identified, which are important for regulation of cellular redox homeostasis (Xiong et al., 2018). Further, genes associated with the methionine sulfoxide reductase system (*msrA* and *msrB*) were recognized in the genome of *LS-ARS2*, which are reported to protect cellular proteins from damage by ROS-mediated oxidation (Oien and Moskovitz, 2019). These genomic signatures encouraged us to find out the potential antioxidant properties of *LS-ARS2* experimentally.

#### 3.5.1 Cation scavenging ability of *LS-ARS2* by ABTS assay

In the ABTS cation radical scavenging assay, the intact cells of *LS-ARS2* showed a 62.14% ABTS scavenging rate, and the heat-lysed cells showed a 71% scavenging rate. Whereas the ABTS scavenging rate of reference strain *L. acidophilus* DDS1 was 47.59% for intact cells and 57.10% for heat-lysed cells (**Figure 4A**). Therefore, the ABTS scavenging rate of *LS-ARS2* was found to be significant, even higher (*p<0.05) than the reference probiotic strain *L. acidophilus* DDS1 for both intact and heat-lysed cells.

**FIGURE 4.**
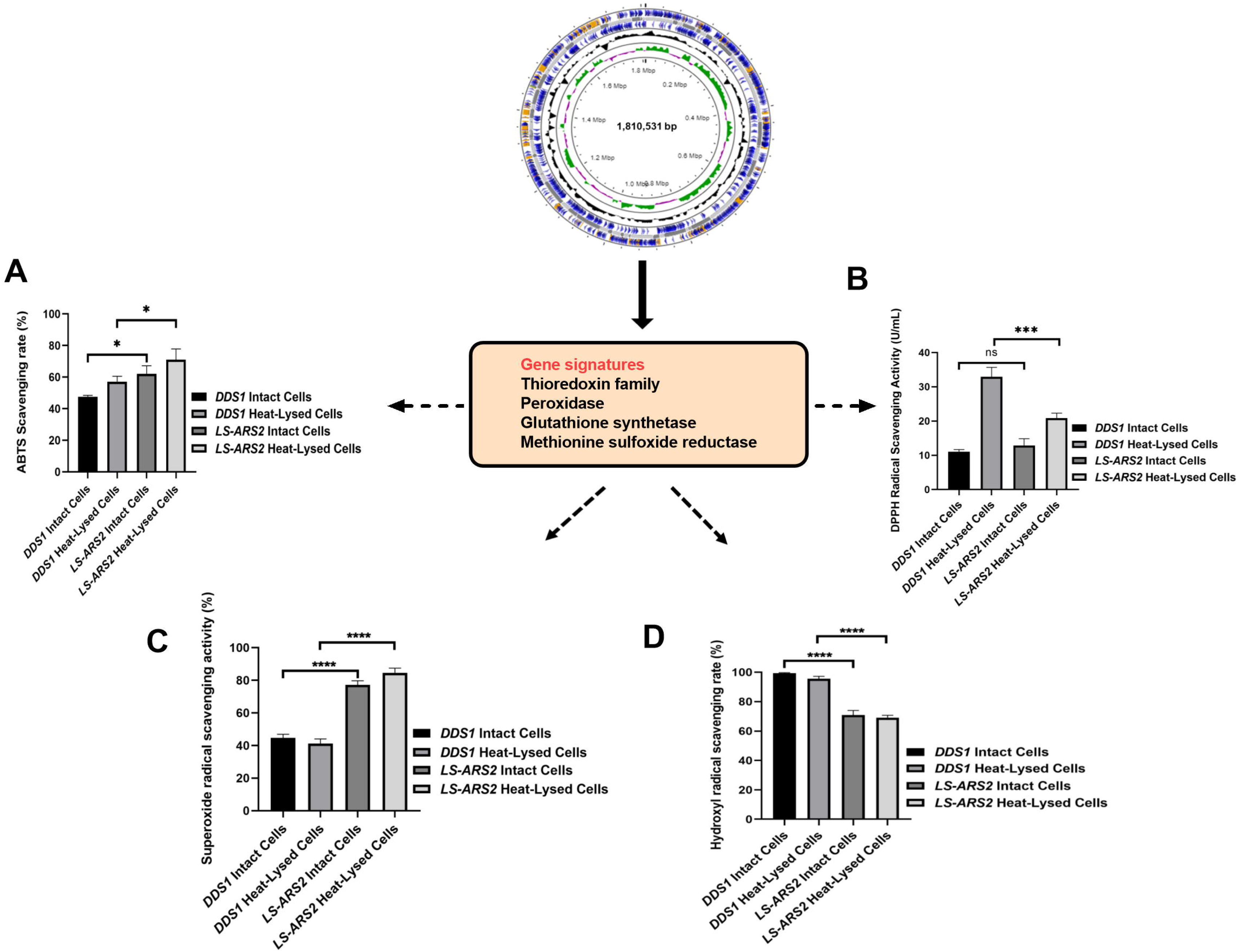
Anti-oxidant properties of *LS-ARS2*. **(A) ABTS cation radical scavenging assay**. The result depicted a significant cation radical scavenging rate of intact and heat-lysed cells of *LS-ARS2*. **(B) DPPH free radical scavenging assay.** The result demonstrated the free radical scavenging activity of *LS-ARS2* intact and heat-lysed cells. **(C) Superoxide free radical scavenging assay.** The data showed the superoxide anion scavenging capacity of *LS-ARS2*. **(D) Hydroxyl free radical scavenging assay.** The result indicated hydroxyl radical scavenging ability of *LS-ARS2* intact and heat-lysed cells. All the antioxidant activities were compared with the clinically used probiotic *L. acidophilus* DDS1. The results were shown as mean ± SD from three independent biological replicates. Statistical analysis was done using one-way ANOVA with a Bonferroni post hoc test. *p<0.05; ****p<0.0001. Consequently, **WGS analysis** showed the *LS-ARS2* genome consists of genes of thioredoxin system as well as glutathione synthetase, which could be potentially associated with the antioxidant properties of *LS-ARS2*.

#### 3.5.2 Free radical scavenging capacity of *LS-ARS2* by DPPH assay

DPPH is used extensively to estimate the free radical scavenging efficacy of antioxidant substances. The transfer of an electron or hydrogen atom to DPPH radical causes a decrease in the absorbance at 517 nm proportionally to the increase of non-radical DPPH (yellow) (Gao et al., 2013). DPPH free radical scavenging activity of *LS-ARS2* whole cell was found to be 12.8 U/mL, which was comparable (p>0.05) to reference probiotic strain *L. acidophilus* DDS1 whole cell (11.08 U/mL). However, the antioxidant activity of *LS-ARS2* heat-lysed cells was found to be significant (20.8 U/mL) but less than the reference probiotic strain *L. acidophilus* DDS1 (32.9 U/mL) (***p<0.0001) (**Figure 4B**). It is to be noted that intact cells of bacteria show the functional properties of the whole cells, whereas heat-lysed form reveals the functionality of intracellular components (Piqué et al., 2019). The intracellular components are strain-specific properties of the probiotics. Therefore, less free-radical scavenging activity of *LS-ARS2* heat-lysed cells than that of DDS1 indicated more significant antioxidant activity of intracellular components of DDS1 than *LS-ARS2*. However, the free radical scavenging capacity of *LS-ARS2* whole cells appeared to be comparable with that of the reference strain, indicating the potential use of *LS-ARS2* as a promising antioxidant agent.

#### 3.5.3 Superoxide radical scavenging ability of *LS-ARS2*

Superoxide anions serve as precursors of singlet oxygen, which indirectly initiates lipid oxidation. The superoxide anion scavenging rate of intact cells of *LS-ARS2* was 77.24%, and heat-lysed cells was 84.61%. Whereas the scavenging rate of reference probiotic strain *L. acidophilus* DDS1 intact cells was 44.68% and heat-lysed cells was 41.15% (**Figure 4C**). Hence, *LS-ARS2* depicted remarkable superoxide anion scavenging potency, even higher (****p<0.0001) than the reference probiotic strain *L. acidophilus* DDS1 for both intact and heat-lysed cells.

#### 3.5.4 Hydroxyl radical scavenging capacity of *LS-ARS2*

Hydrogen peroxide generates hydroxyl radicals in the body as a by-product of various biological processes. This toxic radical initiates indirect oxidation of lipids or proteins and subsequent damage to tissues (Gao et al., 2013). Probiotics may scavenge the radical and reduce such tissue damage. The hydroxyl radical scavenging rate of *LS-ARS2* intact cells was 70.83%, and heat-lysed cells was 69.09%. Reference probiotic strain *L. acidophilus* DDS1 exhibited a 99.41% hydroxyl radical scavenging rate for intact cells and 95.58% for heat-lysed cells. Therefore, the hydroxyl radical scavenging rate of *LS-ARS2* was moderate for both intact cells and heat-lysed cells, although less than the reference probiotic strain *L. acidophilus* DDS1 (****p<0.0001) (**Figure 4D**). However, functional properties like hydroxyl radical scavenging capacity are distinct and independent attributes of probiotics. Less activity of *LS-ARS2* than the reference strain could indicate the limitation of the use of *LS-ARS2* as a hydroxyl radical scavenging agent.

### 3.6 Functional attributes of *LS-ARS2:* Promising antimicrobial properties

#### 3.6.1 Co-culture assay indicated significant anti-*Salmonella* effect of the strain *LS-ARS2*

*S. typhimurium* (ST) is one of the major food pathogens that trigger gut inflammation and life-threatening diarrheal diseases. Therefore, we wanted to evaluate the antimicrobial activity of *LS-ARS2* against *ST*. While characterizing the CFS of *LS-ARS2*, the heat-denatured CFS indicated a zone of inhibition (13.6 mm ± 0.5) similar to the untreated CFS (13.6 mm ± 0.5) in the agar well diffusion assay, whereas the inhibitory effect decreased (0 mm) significantly in the pH-neutralized CFS. This result indicated that the acidic nature of CFS could be the crucial factor for the antibacterial effects of *LS-ARS2* (**Figure 5A**). This result was similar to the reference probiotic strain *L. rhamnosus* GG (*LR-GG*) (p>0.05). The broth microdilution method showed that 10% of the CFS was enough to inhibit the visible growth of the pathogen and was considered the MIP (**Figure 5B**). Agar spot assay also suggested that 10% CFS depicted a bacteriostatic effect, whereas 20% CFS was required for a bactericidal effect (**Figure 5C**). The result shown by *LR-GG* was comparable to *LS-ARS2* (**Figure 5B-5G**). When *LS-ARS2* was co-cultured with *ST*, it effectively hindered the viability of the pathogen with time. A significant (****p<0.0001) reduction in the viable count of the pathogen was observed after 12 h of co-culture, and complete growth inhibition was observed after 24 h (**Figure 6A**).

**FIGURE 5.**
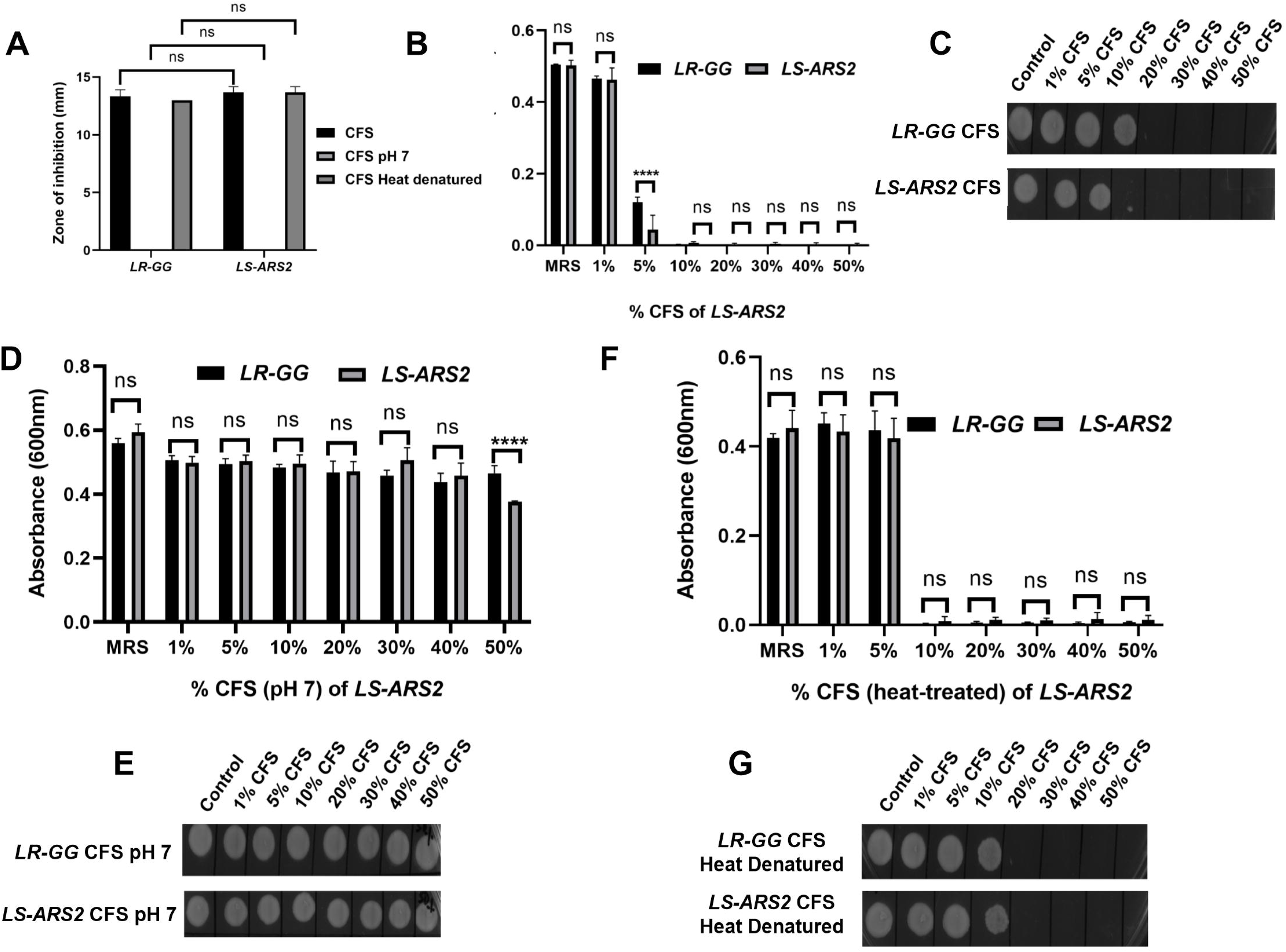
Antimicrobial properties of *LS-ARS2*. **(A) Agar well diffusion assay.** The results showed the effect of normal, pH-neutralized, and heat-denatured cell-free supernatant (CFS) of *LS-ARS2* and *LR-GG* (positive control) on the growth of *S. typhimurium* (ST). The results were shown as mean ± SD based on three independent biological replicates. Statistical analysis was done using two-way ANOVA with Bonferroni post hoc test *p<0.05; ****p<0.0001. **(B, D, F) Determination of the minimum inhibitory percentage (MIP) of *LS-ARS2* CSF for ST.** ST was incubated in various percentages of **(B)** normal, **(D)** pH-neutralized, and **(F)** heat-denatured CFS of *LS-ARS2* and *LR-GG* to estimate the MIP of the CFS against the pathogen. The data presented from three biological replicates; *p<0.05; ****p<0.0001 using two-way ANOVA with Bonferroni post hoc test. **(C, E, G) Estimation of viability of ST in presence of *LS-ARS2* CSF.** From every well of ST grown in *LS-ARS2* CFS and the probiotic *LR-GG* CFS, culture was spotted to find out the bactericidal or bacteriostatic effect (**C**, **E**, **G**). All the data are represented as the mean ± SD from three independent biological replicates.

**FIGURE 6.**
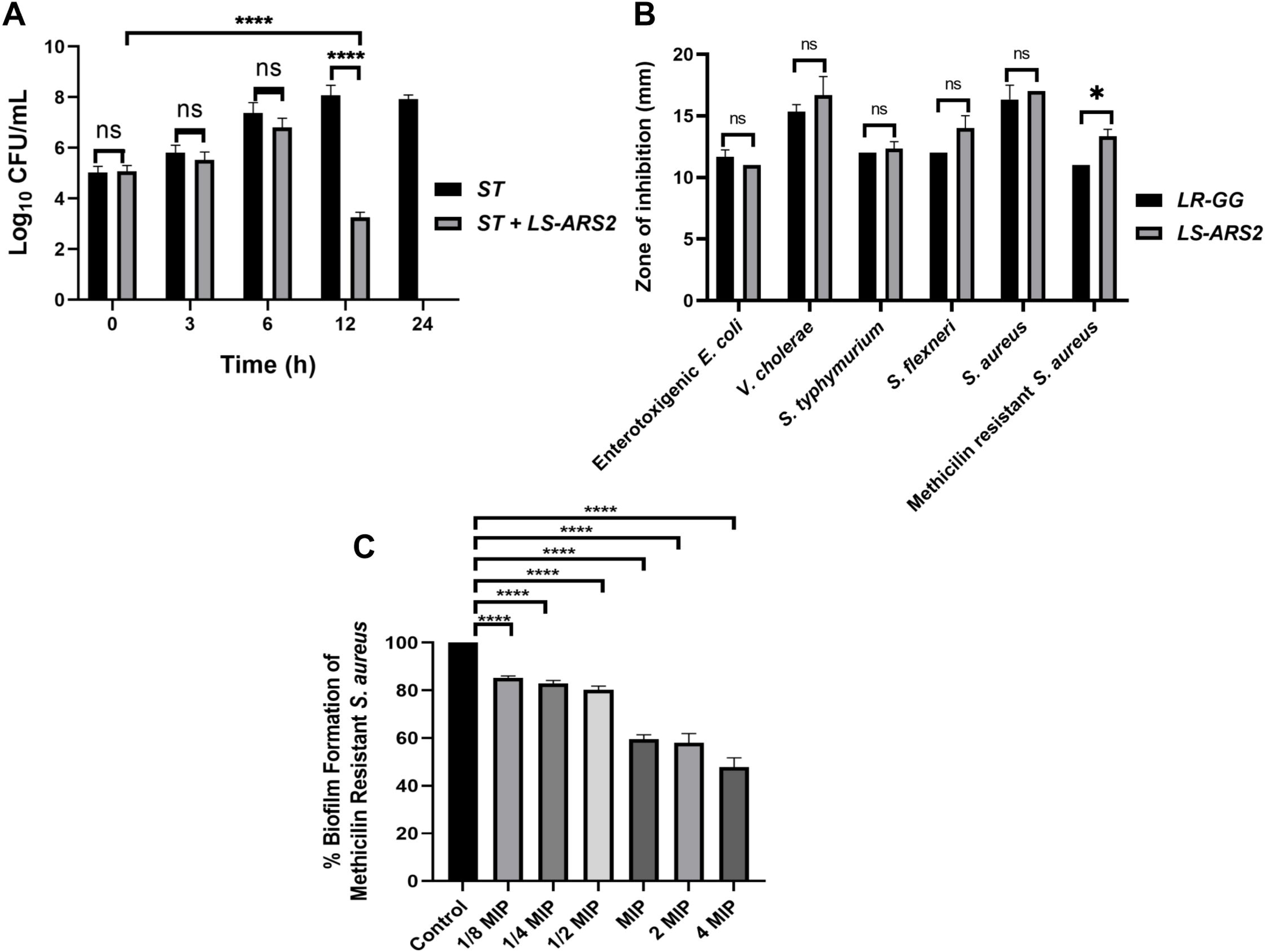
Antimicrobial properties of *LS-ARS2*. **(A) Co-culture assay.** Co-culture of *S. typhimurium* (ST) with *LS-ARS2* at multiple time periods illustrated the *LS-ARS2* cytotoxic effect to the pathogen. After 12 h of co-culture, a significant reduction (****p<0.0001) in the viable cell number of *S. typhimurium* was observed. Whereas, after 24 h of culture, the viable cell number of ST was decreased to zero (data presented from three biological replicates with multiple technical repeats; *p<0.05; ****p<0.0001 using a two-way ANOVA with a Bonferroni post hoc test). **(B) Agar well diffusion assay**. The result showed diverse inhibition zones, signifying the antimicrobial activity of *LS-ARS2* and *LR-GG* (positive control) against multiple Gram-positive and Gram-negative enteric pathogens. The data were represented as mean ± SD based on three independent biological replicates. Statistical analysis was done using two-way ANOVA with Bonferroni post hoc test; p>0.05, *p<0.05. **(C) Anti-biofilm Assay.** Cell-free supernatant (CFS) of *LS-ARS2* showed a significant reduction in the biofilm formation of Methicillin-resistant *Staphylococcus aureus* ATCC 700699. The results were represented as the mean ± SD based on three independent biological replicates. Statistical analysis was done using one-way ANOVA with a Bonferroni post hoc test; ****p<0.0001.

#### 3.6.2 Robust antimicrobial potential of *LS-ARS2* across the diverse pathogen

It is desirable for a potential probiotic to possess robust antimicrobial potential for a wide range of pathogens. Indeed, *LS-ARS2* demonstrated remarkable inhibitory activity against all studied Gram-positive and Gram-negative enteric pathogens (**Figure 6B**). Among them, the strain showed the highest antimicrobial activity for *S. aureus*, *V. cholerae*, and *S. flexneri*. It should be emphasized that *V. cholerae* and *S. flexneri* were multi-drug-resistant clinical isolates. Moreover, the strain also showed a significant inhibitory effect against Gram-positive methicillin-resistant *S. aureus*. The antimicrobial activity of *LR-GG* was comparable to *LS-ARS2* (**Figure 6B**) (p>0.05).

#### 3.6.3 Anti-biofilm effect of *LS-*ARS2 for MRSA

The pathogenic biofilms pose serious hazards in the food and clinical industries, affecting public health. Therefore, safe applications like probiotics are encouraged to reduce the prevalence of the growth of pathogenic biofilms. MRSA is known to be a strong biofilm-former, and MRSA-biofilm is a part of their virulence factor (Kaushik et al., 2024).

To estimate the anti-biofilm effect of *LS-ARS2*, first the MIP of CFS for MRSA was determined (10% of *LS-ARS2-*CFS, **Supplementary Figure S2**). Further, 1/8 MIP to 4 MIP (1.25%-40%) of *LS-ARS2*-derived CFS were used to study the formation of MRSA-biofilm. Our result indicated a significant reduction (****p<0.0001) in the formation of MRSA-biofilm at all studied concentrations. Moreover, *LS-ARS2*-derived CFS was able to reduce biofilm formation even at 1.25%. These results illustrated the potential of the strain to mitigate the formation of pathogenic biofilm (**Figure 6C**).

#### 3.6.4 Prediction of bacteriocin in the *LS-ARS2* genome

The anti-*Salmonella* effect of *LS-ARS2* was eliminated in the CFS, in which the pH was neutralized, indicating the presence of pH-sensitive bacteriocins. This result motivated us to further dig-in to the *LS-ARS2* genome in search of bacteriocin-encoding gene signatures. It was extremely exciting to find that one area of interest (AOI) region, responsible for BCG, was identified in the genome of *LS-ARS2* (**Figure 7**). The BCG was located within contig 32.1 of salivaricin_P_chain_b class. The AOI of contig 32.1 resides with salivaricin_P_chain_b (bit score = 134.806) and salivaricin_P_chain_a (bit score = 122.479) (Barrett et al., 2007). Besides, *LS-ARS2* contains various ORFs encoding miscellaneous bacteriocins (Tenea and Ascanta, 2022), accessory transport, and sensor proteins as indicated in **Figure 7** (Tenea and Ascanta, 2022).

**FIGURE 7.**
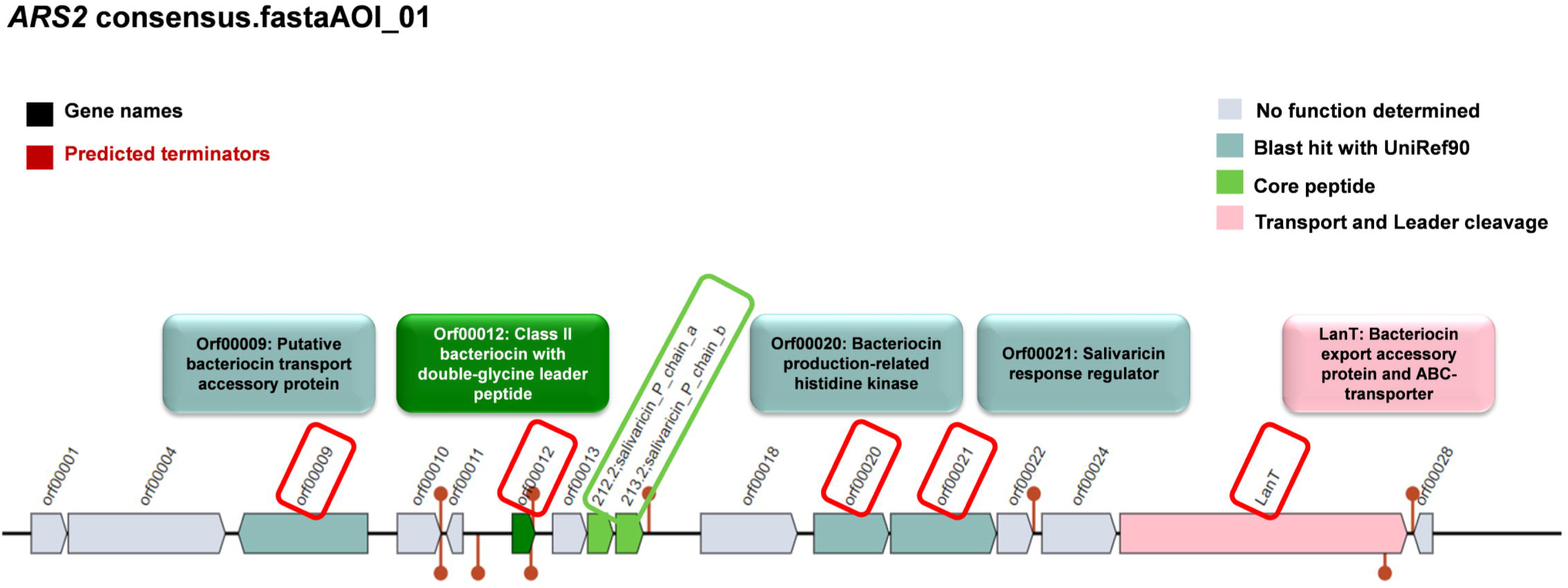
Prediction of genes for bacteriocin and accessory proteins in Biosynthetic Gene Clusters (BCGs) organization in the genome of *LS-ARS2* (BAGEL4 analysis).

### 3.8 Functional attributes of *LS-ARS2:* Prediction from genomic analyses

#### 3.8.1 Association of beneficial metabolite-producing pathways in the *LS-ARS2* genome

Functional annotation of the *LS-ARS2* genome showed 30 enriched KEGG pathways. Among them, metabolic pathways contained the highest gene count (approximately 40), followed by the pathway responsible for biosynthesis of secondary metabolites (approximately 20). Further, the KEGG pathway related to microbial metabolism in diverse environments with an approximate gene count of 10 indicated the adaptation capability of *LS-ARS2* in various niches. Moreover, the strain contained numerous genes involved in major metabolic pathways like amino acid biosynthesis and carbon metabolism. Interestingly, abundant genes associated with essential amino acid metabolism (methionine, threonine, phenylalanine, and tryptophan) were detected in the *LS-ARS2* genome, which indicated the potential health-benefits of the strain (**Figure 8A**).

**FIGURE 8.**
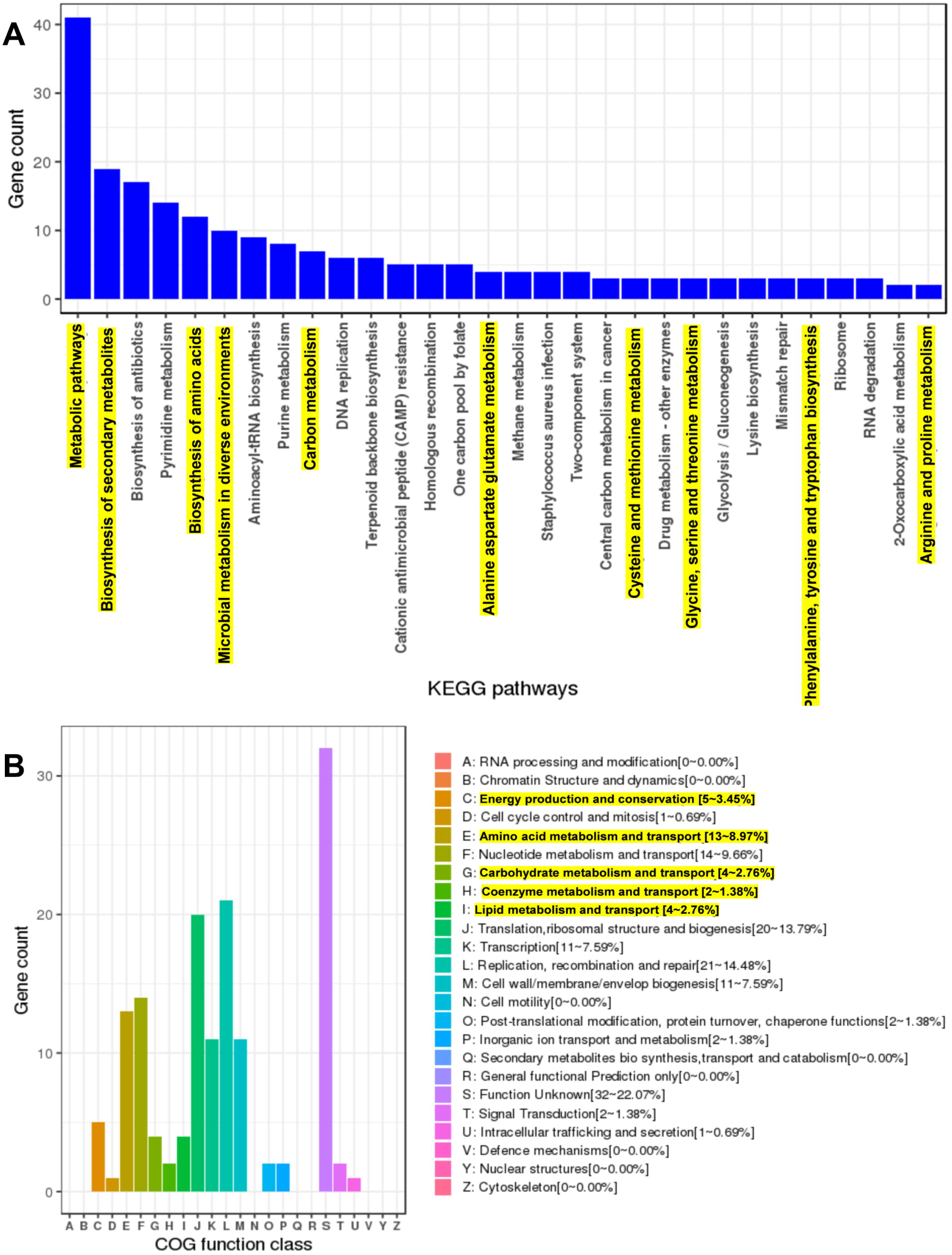
Genome mining of the *LS-ARS2* genome and prediction of major pathways and COG categories. **(A) The KEGG enriched pathways.** Top 30 KEGG enriched pathways were analysed from the genome of *LS-ARS2*. Metabolic pathways, biosynthesis of secondary metabolites, and biosynthesis of antibiotics were three major pathways. Highlighted KEGG pathways showed the gene count of significant metabolite-producing pathways. **(B) WGS analysis identified major COG categories in the *LS-ARS2* genome.** Highlighted COG categories showed the abundant gene count responsible for the production of beneficial metabolites.

Next, the protein-coding genes of the *LS-ARS2* genome were found to be 1586, among which 1527 (96.28%) genes were classified into diverse COG functions. The annotated genes were found to be involved in several physiological processes. Functional annotation revealed the association of significant COG categories like energy production and conversion (5-3.45%), amino acid metabolism and transport (13-8.97%), carbohydrate metabolism and transport (4-2.76%), coenzyme metabolism and transport (2-1.38%), and lipid metabolism and transport (4-2.76%) (**Figure 8B**). Corroborating with the KEGG pathways, the COG analysis also showed the abundance of the genes that could be responsible for the production of beneficial metabolites (amino acids, vitamins) and amino acid or carbohydrate metabolism. The enrichment of metabolic genes in the *LS-ARS2* genome further strengthened the potential of the strain to survive in diverse environments.

#### 3.8.2 Detection of carbohydrate active enzymes (CAZymes)

Carbohydrate active enzymes (CAZymes) enable bacteria to adapt in harsh environments by utilizing complex carbohydrate molecules (Liang et al., 2024). KEGG pathways and COG analysis revealed the abundance of genes associated with carbohydrate metabolism and transport, which could enable microbial metabolism in diverse environments. Further mining into the *LS-ARS2* genome revealed the presence of CAZymes that belonged to the glycoside hydrolase family (GH), glycosyltransferases family (GTs), and carbohydrate binding modules (CBM) (**Table 4**). CAZymes of the GH family are responsible for the hydrolysis of complex carbohydrates. In the *LS-ARS2* genome, GH13, GH31, GH77, GH32, GH65, and GH37 were detected. Among these, GH31, GH32, and GH65 are reported to participate in the metabolism of monosaccharides and oligosaccharides such as glucose, galactose, fructose, xylose, or arabinose (Kavanova et al., 2024). Whereas CAZyme families GH13 and GH77 are involved in the hydrolysis of polysaccharides such as α-glucans, β-glucans, or arabinoxylans. The GH37 family is known to be associated with the degradation of trehalose (Shrestha et al., 2024). In addition, CAZymes involved in the transfer of sugar moieties are termed glycosyltransferases (GTs), which play a key role to form surface structures sensed by the host immune system (Kandasamy et al., 2024). GTs annotated in the *LS-ARS2* genome were GT1, GT2, GT4, GT5, GT26, GT28, GT35, and GT51. Moreover, two types of carbohydrate binding modules (CBM), CBM48 and CBM50, were detected in the *LS-ARS2* genome. CBM helps in the binding of some carbohydrate hydrolases to augment the hydrolysis of polysaccharides. Functional annotation also revealed the presence of few sugar transporter genes (*pts13C, manL, dhaM, srlB, mtlA*) in the *LS-ARS2* genome, indicating the strong carbon transporting ability of the strain, which is important to inhabit at the gastrointestinal environment as well as other ecological niches (Kandasamy et al., 2024) (**Supplementary Table S2**).

**Table 4.**
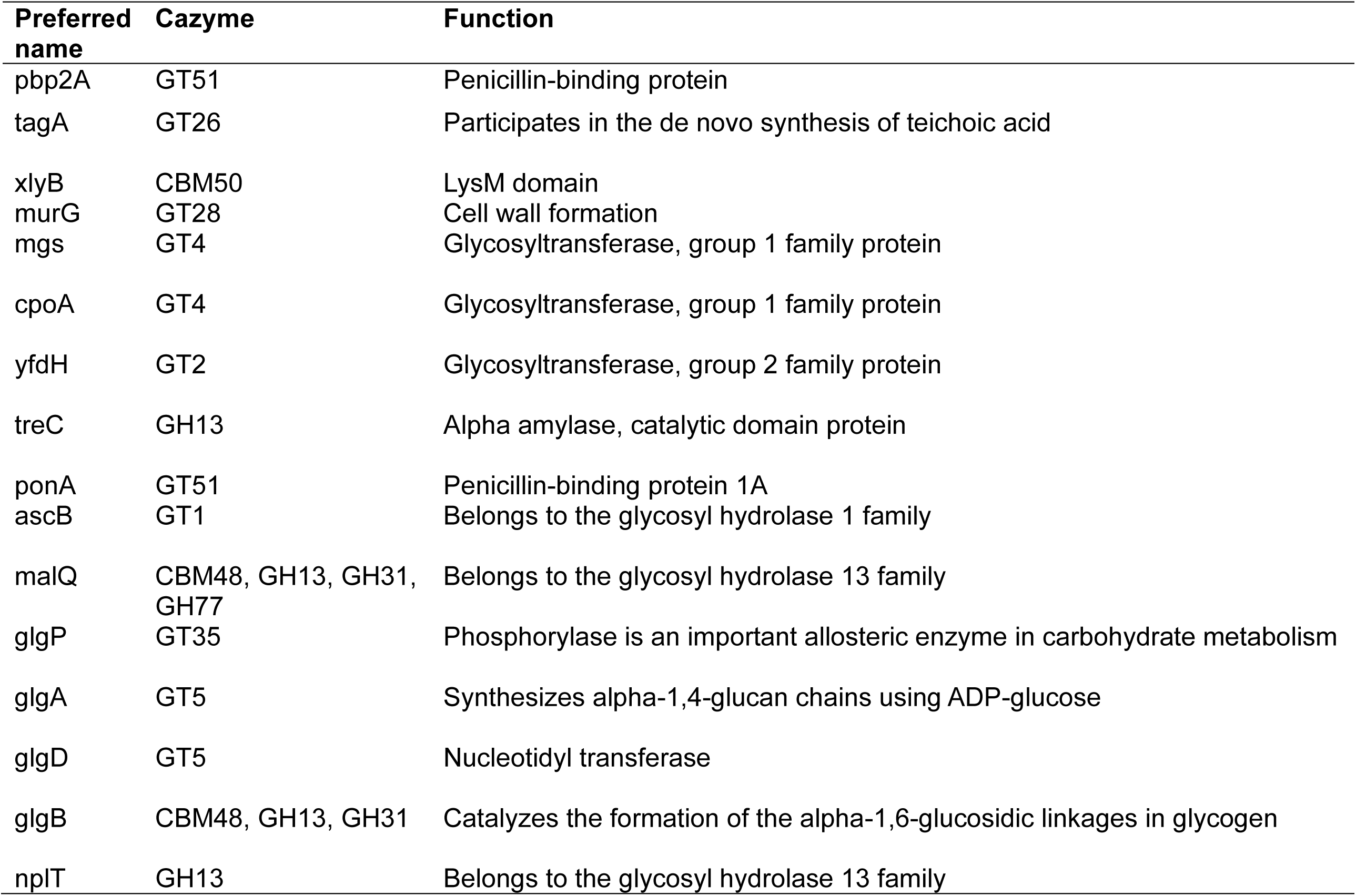
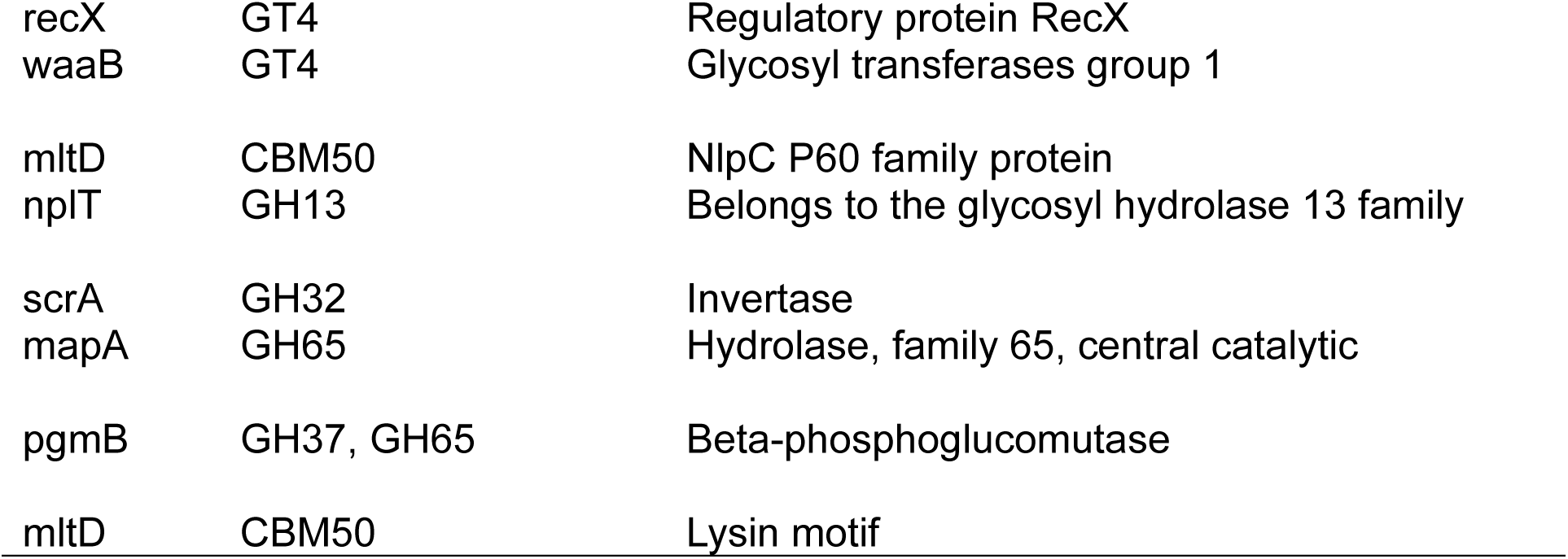
Carbohydrate active enzymes (CAZymes) detected in the *LS-ARS2* genome.

| Preferred name | Cazyme | Function |
| --- | --- | --- |
| pbp2A | GT51 | Penicillin-binding protein |
| tagA | GT26 | Participates in the de novo synthesis of teichoic acid |
| xlyB | CBM50 | LysM domain |
| murG | GT28 | Cell wall formation |
| mgs | GT4 | Glycosyltransferase, group 1 family protein |
| cpoA | GT4 | Glycosyltransferase, group 1 family protein |
| yfdH | GT2 | Glycosyltransferase, group 2 family protein |
| treC | GH13 | Alpha amylase, catalytic domain protein |
| ponA | GT51 | Penicillin-binding protein 1A |
| ascB | GT1 | Belongs to the glycosyl hydrolase 1 family |
| malQ | CBM48, GH13, GH31, GH77 | Belongs to the glycosyl hydrolase 13 family |
| glgP | GT35 | Phosphorylase is an important allosteric enzyme in carbohydrate metabolism |
| glgA | GT5 | Synthesizes alpha-1,4-glucan chains using ADP-glucose |
| glgD | GT5 | Nucleotidyl transferase |
| glgB | CBM48, GH13, GH31 | Catalyzes the formation of the alpha-1,6-glucosidic linkages in glycogen |
| npIT | GH13 | Belongs to the glycosyl hydrolase 13 family |
| recX | GT4 | Regulatory protein RecX |
| waaB | GT4 | Glycosyl transferases group 1 |
| mltD | CBM50 | NlpC P60 family protein |
| nplT | GH13 | Belongs to the glycosyl hydrolase 13 family |
| scrA | GH32 | Invertase |
| mapA | GH65 | Hydrolase, family 65, central catalytic |
| pgmB | GH37, GH65 | Beta-phosphoglucomutase |
| mltD | CBM50 | Lysin motif |

### 3. 9 Metabolomics of *LS-ARS2:* Genomic prediction coupled with experimental validation by HRMS analysis

#### 3.9.1 Prediction of primary and secondary metabolites in the *LS-ARS2* genome

Microorganisms produce various substances (primary and secondary metabolites) that modulate host metabolism and immunity. Metabolic gene cluster (MGC) is defined as a cluster of genes that encodes different enzymes of the same metabolic pathway. The gutSMASH tool predicts genes involved in bioenergetics and metabolism in anaerobic bacteria (Pascal Andreu et al., 2021). In our study, gutSMASH found one MGC region, pyruvate to acetate-formate type (region 58.1), in the *LS-ARS2* genome (**Figure 9A**). This region encodes for short-chain fatty acids (SCFAs), primarily acetate, butyrate, and propionate, which could have a positive impact on gut-health (Donia and Fischbach, 2015). The KnownClusterBlast output indicated high sequence similarity between the predicted *LS-ARS2* MGC and other *Lactobacillus* strains (**Figure 9B**). Interestingly, among all the *Lactobacillus* strains, the highest gene similarity of the Pyruvate2acetate-formate region was found with other *L. salivarius* strains with the same region and known function. The KnownClusterBlast output thus strongly supports the potential production of primary metabolites (SCFAs) from *LS-ARS2*. The gutSMASH results nicely corroborate with the previous KEGG pathways and COG analysis, where abundant gene count was detected for metabolic pathways (specifically carbohydrate metabolism and transport) and energy production.

**FIGURE 9.**
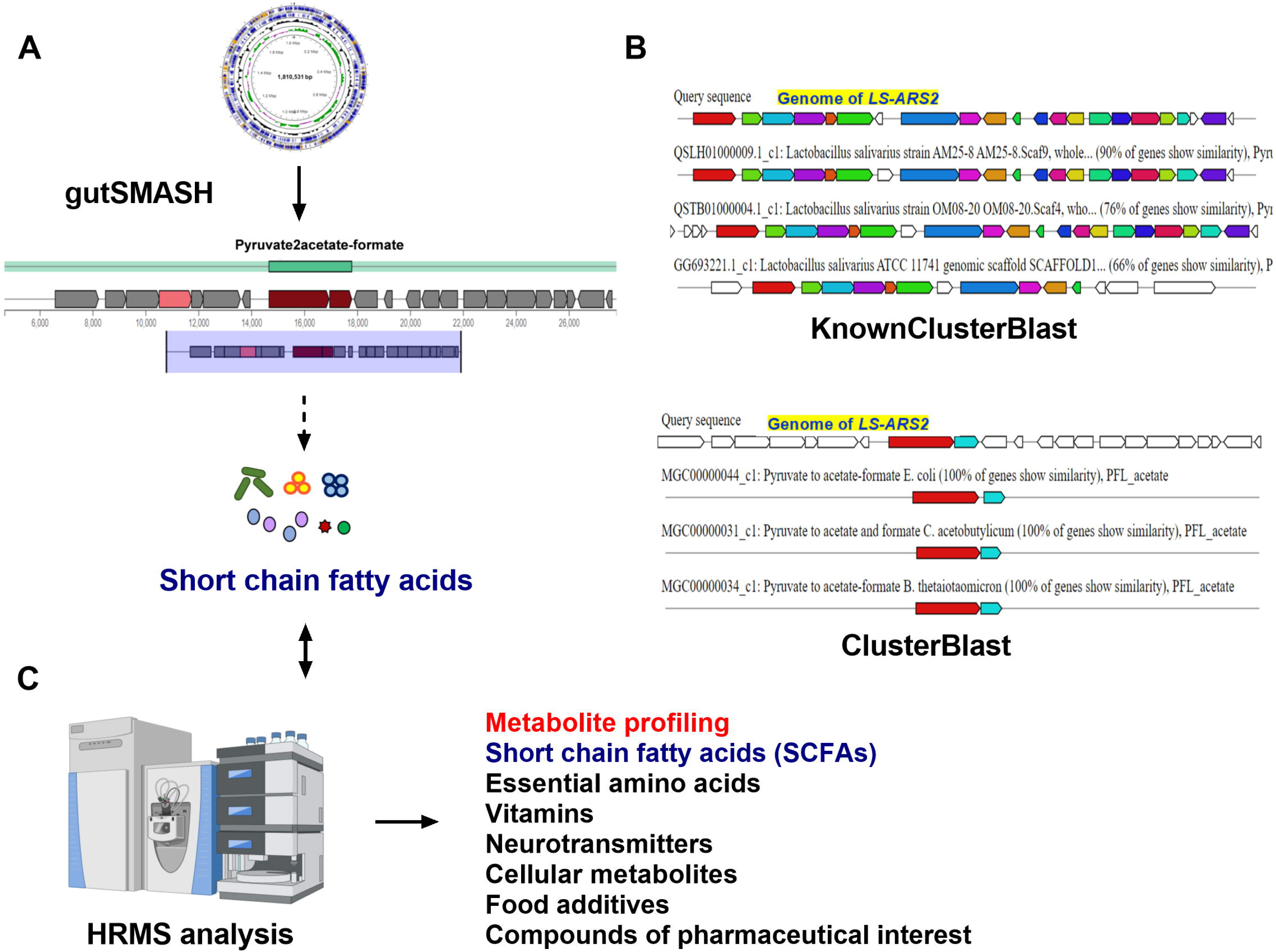
Prediction of genes for primary metabolites in Metabolic Gene Clusters (MCGs) of the *LS-ARS2* genome. **(A) GutSMASH run of the *LS-ARS2* genome.** GutSMASH predicted one MGC region, type pyruvate to acetate-formate, which is responsible for the production of SCFAs. **(B) Gene Cluster Comparative Analysis.** Comparative analysis between different bacterial reference genes (pre-computed in gutSMASH) for the most similar metabolic gene clusters (MGCs) based on a gutSMASH run. Genes with the same colour indicated putative homologs based on significant Blast hits between *LS-ARS2* and the reference bacterial genes. The **KnownClusterBlast analysis** showed the gene similarity of the predicted region (Pyruvate2acetate-formate) with MGCs associated with known functions of other *Lactobacillus* reference genomes. The **ClusterBlast analysis** output showed that the predicted metabolic gene cluster (MGC) (Pyruvate2acetate-formate) does not have homologous MGCs among other *Lactobacillus*. White genes have no relationship. (C) Detection of beneficial metabolites in the *LS-ARS2* CFS using HRMS analysis (also see **Table 5** for details).

**Table 5.**
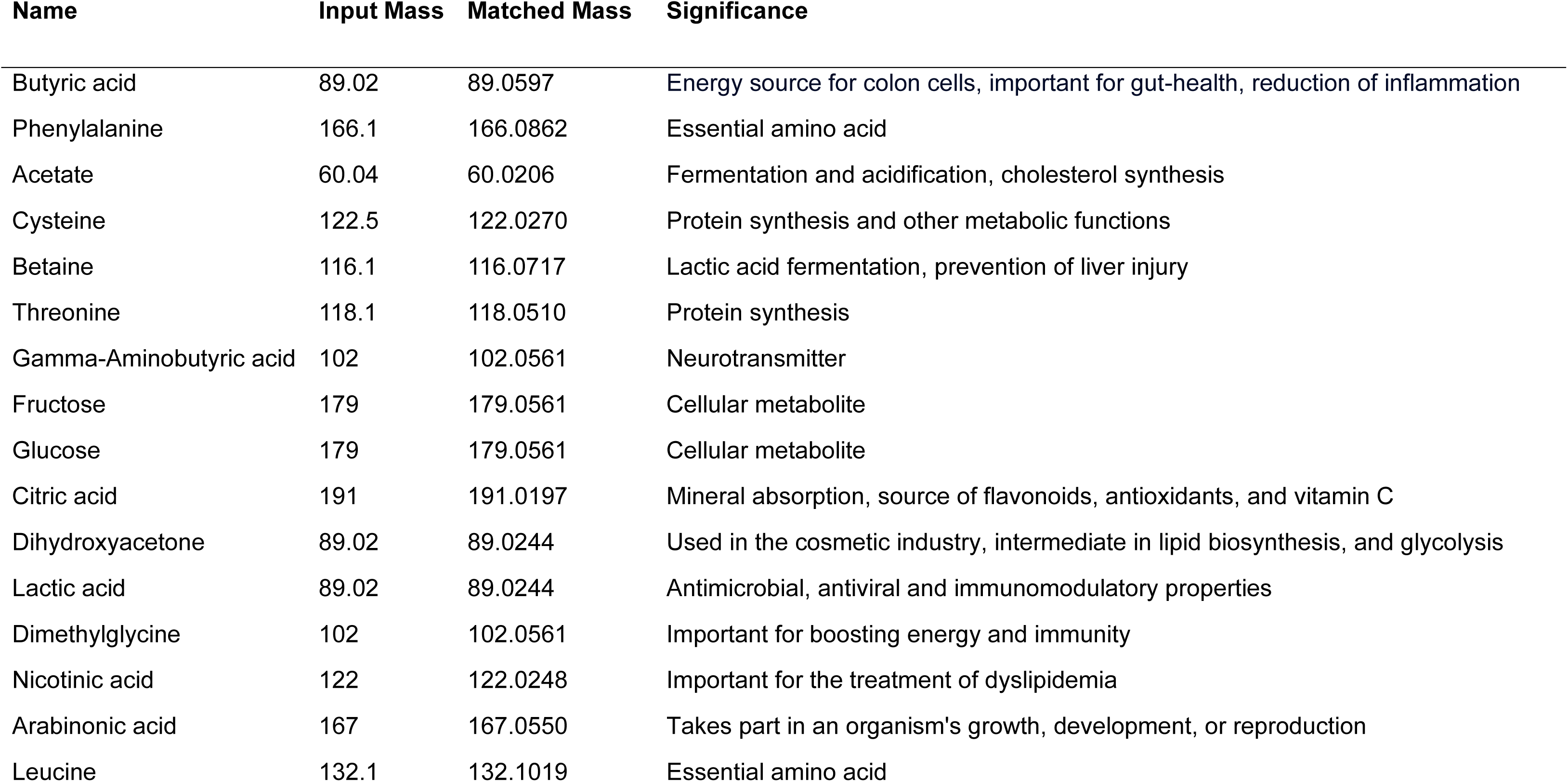
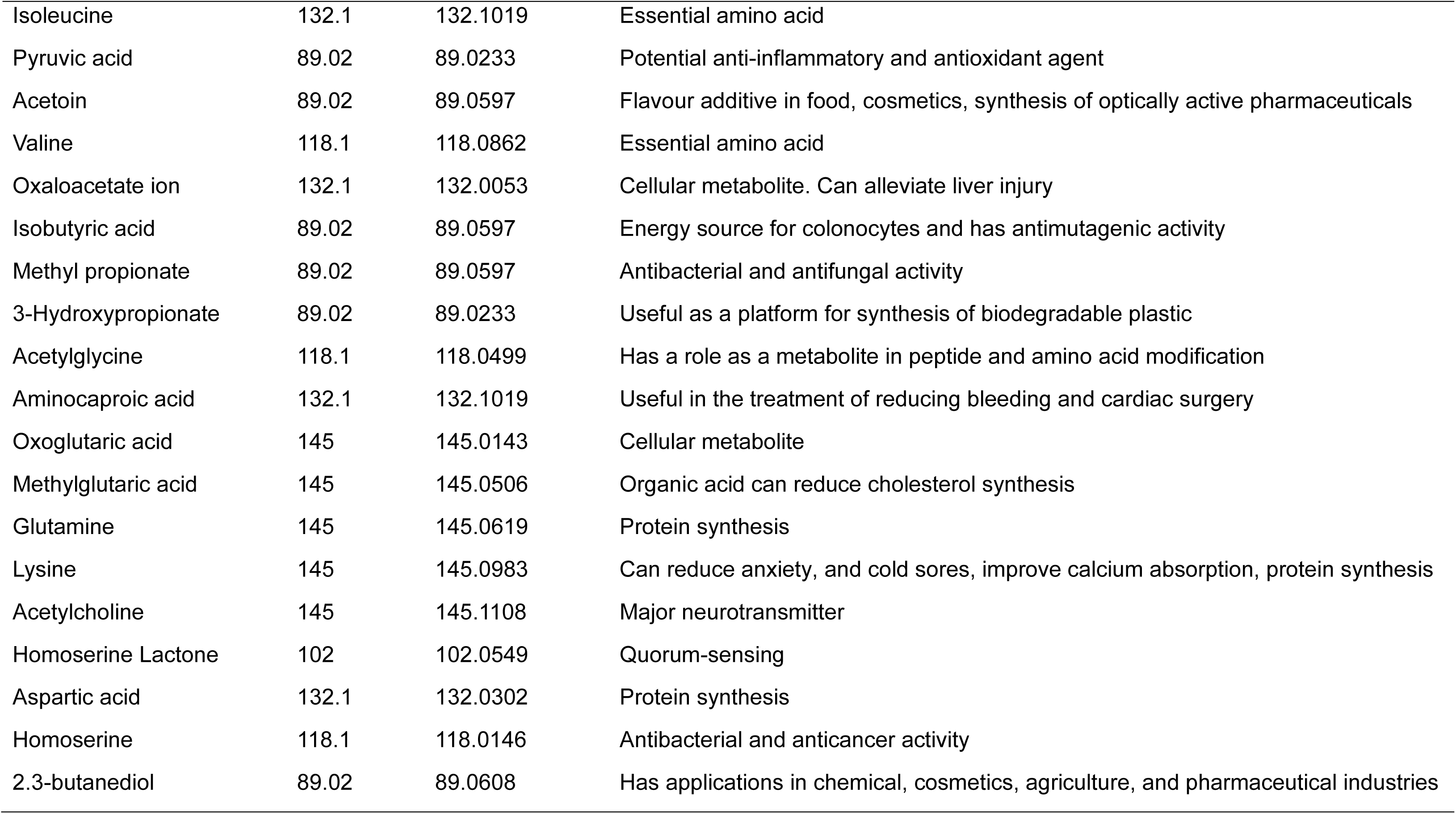
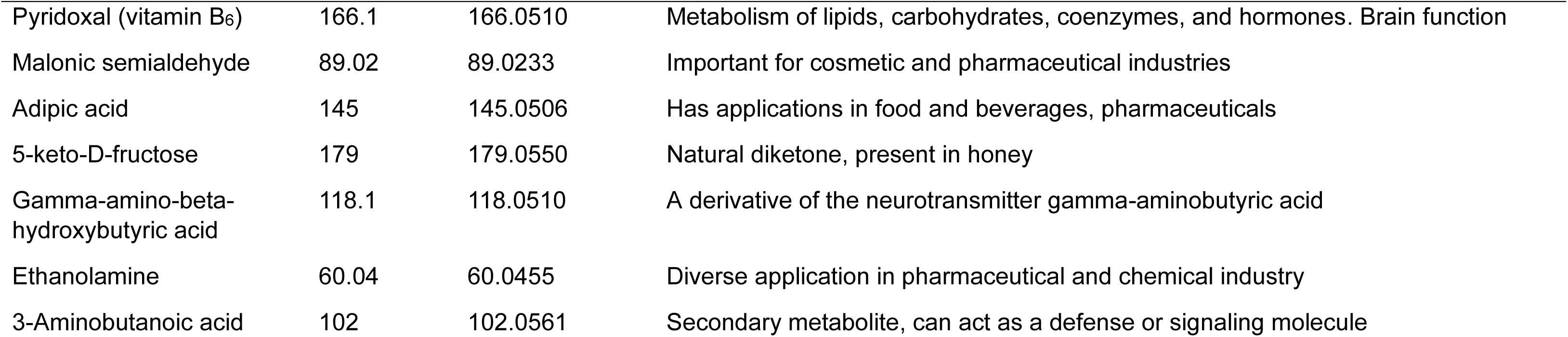
Metabolite Profiling of the Cell-Free Supernatant (CFS) of *LS-ARS2*.

The KEGG pathway analysis indicated the richness of genes responsible for the biosynthesis of secondary metabolites, which are vital for the antibacterial, antifungal, and antitumor activities of the microorganisms (Tenea and Ascanta, 2022). The web server, antiSMASH, predicts NRPs, PKs, terpenes, and RiPP-like peptides with various known or putative secondary metabolites in the genome of bacteria. Fascinatingly, further genome mining of the *LS-ARS2* genome by antiSMASH predicted one secondary metabolite-producing region (region 39.1) named T3PKS region (T3PKS: Chal_sti_synt_N, Chalcone and Stilbene synthases, N-terminal domain) within the *LS-ARS2* genome (**Supplementary Table S6)**. Polyketide synthases (PKSs) are related to the synthesis of antibiotics and pharmaceutical products with various roles in food processing (Naughton et al., 2017). Microbial Type III PKSs (T3PKS) are involved in the biosynthesis of some lipidic compounds and various secondary metabolites with significant biological functions (Katsuyama and Ohnishi, 2012).

#### 3.9.2 Beneficial metabolites in *LS-ARS2* CFS: HRMS analyses

The detailed genome mining and functional annotation of the *LS-ARS2* genome showed the abundance of the genes responsible for synthesizing beneficial metabolites (**Figure 8A-B and 9A-B**). Next, we were excited to explore the metabolite profiling of the strain. Our HRMS study indicated a wide range of antibacterial metabolites, and organic acids (for example, lactic acid, acetic acid, butyric acid, and homoserine) were present in the CFS of *LS-ARS2*. Several SCFAs, like butyric acid, isobutyric acid, lactic acid, acetic acid, and hydroxypropionic acid, were also prominent in CFS, corroborating output from gutSMASH (**Figure 9C**). These compounds have been shown to function as major antimicrobial components and immunomodulators (Dash et al., 2023). Many essential amino acids like leucine, isoleucine, valine, phenylalanine, threonine, and lysine were prevalent in the CFS (see discussion), which were nicely in support of the findings from KEGG pathways analysis (**Figure 8A**). Vitamins such as pyridoxal (vitamin B_6_) and nicotinic acid (vitamin B_3_) were also found (**Table 5 and Figure 9C**). Therefore, our HRMS analysis beautifully validated the findings from the WGS analysis and confirmed the presence of beneficial metabolites in the CFS of *LS-ARS2*.

## 4. Discussion

Despite their promising potential as probiotics, very few *L. salivarius* strains are available at an industrial scale as commercial products [Guerrero Sanchez et al. 2022]. Our results illustrate that *LS-ARS2* is a biofilm-former LAB, that exhibits significant antioxidant, antibacterial, as well as antibiofilm potential. Metabolic profiling of *LS-ARS2*-CFS indicates the presence of diverse health-promoting metabolites as well as other compounds with various applications as food, poultry, and cosmetic products. Therefore, *LS-ARS2* appears as a promising candidate for diverse lucrative applications, particularly in food and therapeutic industries (see below for details).

First, the WGS analysis together with the *in vitro* assays establishes the probiotic attributes of *LS-ARS2.* The strain could potentially withstand the acidic and alkaline environments of the stomach and small intestine, respectively. Biofilm formation ability, aggregation efficacy, as well as adhesion and colonization potential, could facilitate *LS-ARS2* for an efficient and longer stay in the gut. Moreover, the safety attributes of the strain are also assured, although at the *in vitro* level.

Dysbiotic gut often disrupts the intricate oxidative balance (Jm, 1985). Supraphysiological levels of all the reactive oxygen species (ROS), like hydrogen peroxide and superoxide anions, cause severe tissue damage and organ dysfunction (Barry, 1991). The excess level of ROS triggers chronic inflammation that results in gut-inflammatory diseases (ulcerative colitis, intestinal bowel syndrome, colorectal cancer) as well as diseases related to gut-organ axes (Ren et al., 2023). The significant antioxidant activities of *LS-ARS2* reveal the potential of the strain to reduce oxidative stress in the gut. Interestingly, the functional annotation of the *LS-ARS2* genome beautifully corroborates our *in vitro* investigations **(Figure 4),** indicating that supplementation of *LS-ARS2* could contribute to the oxidative balance in the gut.

Chronic infections by pathogenic biofilms are clinically challenging. MRSA, particularly in biofilm form, is a leading cause of hospital-acquired infections that causes significant morbidity and mortality. MRSA-biofilm facilitates the pathogen to invade, spread, and resist antimicrobial treatments (Ciofu et al., 2022). In this background, it is extremely exciting to find that the CFS of *LS-ARS2* inhibits MRSA-biofilm. Application of *LS-ARS2* could therefore provide a green therapeutic window for combating pathogenic-biofilm.

WGS analysis predicts the presence of the MCG region (encodes SCFAs) in the genome of *LS-ARS2*. The HRMS study indicates the presence of diverse SCFAs (acetate, propionate, butyrate, lactic acid, etc.) in the CFS of *LS-ARS2*. Thus, genomic characterization coupled with metabolomic studies (**Figure 9**) confirms that, like other probiotic bacteria, SCFAs in the CFS of *LS-ARS2* could offer beneficial roles to the host-gut. These SCFAs eliminate pathogens by entering through their cell membrane and acidification of their alkaline cytoplasm. Further, SCFAs have numerous health-promoting properties, including antibacterial, antiviral, immunomodulatory, and anticancer effects (Dash et al., 2023). Specifically, butyrate is an energy source for colonocytes (Pedroza Matute and Iyavoo, 2023). Therefore, intake of *LS-ARS2* could facilitate maintaining gut-health.

Bacteriocins, produced by LAB, are ribosomally synthesized antimicrobial peptides. BAGEL4 analysis of the *LS-ARS2* genome shows the presence of one class II bacteriocin-producing BCG (salivaricin_P_chain_b) and bacteriocin accessory as well as export proteins, which supports the probable presence of bacteriocin in the *LS-ARS2* genome. Bacteriocins and associated proteins retain their activity preferably at acidic pH rather than at alkaline or neutral pH (Magnusson and Schnürer, 2001). In the present study, the antimicrobial effect of the *LS-ARS2* CFS against *Salmonella* was not found for pH-neutralized CFS, indicating the possible existence of pH-sensitive bacteriocin-like peptides in the CFS.

HRMS study indicates *LS-ARS2* CFS consists of metabolite betaine, which is an essential osmolyte and neuroprotectant with significant health-improving roles (Arumugam et al., 2021). *LS-ARS2* CFS also consists of gamma-aminobutyric acid (GABA), one of the principal neurotransmitters. GABA significantly regulates nerve cell hyperactivity associated with anxiety, stress, convulsions, epilepsy, Parkinson’s, and Alzheimer’s diseases. Several studies have also reported the immunological and antimicrobial functions of GABA (Bs et al., 2021). Further, genomic analysis reveals the presence of CAZymes in the *LS-ARS2* genome. Bacterial strains use CAZymes for carbohydrate degradation. Therefore, CAZymes in the *LS-ARS2* not only could participate in utilizing complex carbohydrates and thrive in the host intestinal ecosystem; they also could further stimulate the growth of other probiotics by providing the carbon source for them. Therefore, as a nutritional supplement, *LS-ARS2* with CAZymes could facilitate the overall good bacterial distribution and improve gut-health.

Food supplements are added to the diet to maintain the nutritional balance of the body. HRMS analysis indicates *LS-ARS2* CFS is enriched with several beneficial metabolites that highlight the potential of the strain to be used as a dietary supplement. For instance, essential amino acids like threonine, lysine, phenylalanine, leucine, isoleucine, and valine are detected in the CFS of *LS-ARS2.* Next, vitamins nicotinic acid (vitamin B_3_) and pyridoxal (vitamin B_6_) are also found in *LS-ARS2* CFS, indicating that application of the strain could reduce the severity of clinical vitamin deficiencies (Gasperi et al., 2019; Stach et al., 2021). Moreover, *LS-ARS2* CFS is enriched with metabolites like pyruvic acid, oxaloacetate, and oxoglutaric acid, which are reported to have health-promoting effects (Dash et al., 2023; Wernerman et al., 1990).

Besides, *LS-ARS2* CFS is also enriched with metabolites important for food processing and cosmetic industries. Metabolites like citric acid, dihydroxyacetone, and acetoin have extensive applications in the poultry, food, and cosmetic industries (Dash et al., 2023). In our genomic study, antiSMASH predicts the presence of the T3PKS region, which is known to encode substances with enormous functions in food processing industries. Other detected metabolites in *LS-ARS2* CFS, such as adipic acid, can be used as food additives (Horn et al., 1957), whereas 3-Hydroxypropeonate, 2.3-butanediol, and malonic semialdehyde have diverse applications in the pharmaceutical, cosmetics, and agriculture industries (Zhao and Tian, 2021; Gu et al., 2022; Dash et al., 2023).

## 5. Conclusion

We present a full-fledged study on the probiotic attributes of *LS-ARS2*. a) The remarkable anti-oxidant and anti-bacterial potential of *LS-ARS2* encourages further studies of the strain for health-promoting applications. b) Strong biofilm-forming ability of *LS-ARS2* ensures efficient and longer stay in the GI tract. The significant anti-biofilm activity of *LS-ARS2* CFS for MRSA again suggests the use of *LS-ARS2* as a promising antimicrobial agent to inhibit pathogenic biofilm formation. c) As a gut microbe, *LS-ARS2* could naturally re-establish the gut-microbial composition and thereby improve the metabolic status of the gut and prevent dysbiosis. Further, the identification of beneficial metabolites in *LS-ARS2-*CFS with antimicrobial and immunomodulatory properties indicates the therapeutic application of *LS-ARS2* in different diseases. However, *in vivo* studies with gut-inflammatory disease models would reaffirm that.

Potential ability of *LS-ARS2* to stably colonize in the GI tract and other probiotic attributes, including beneficial metabolic profiling of *LS-ARS2-*CFS, indicate the promising application of the strain as a food supplement. Further, myriads of secondary metabolites with industrial applications were detected in *LS-ARS2* CFS, which supports the potential use of the strain in the food as well as poultry and cosmetic industries. Together, the study indicates *LS-ARS2* demands further studies for its potential application in the future. In summary, the study offers a novel and promising probiotic strain, *LS-ARS2,* that demands *in vivo* research, particularly focusing on gut-disease models for their potential applications in the near future.

## Supporting information

pdf attached

## Abbreviations

GI: gastrointestinal
WHO: World Health Organization
WGS: Whole Genome Sequencing
ATCC: American Type Culture Collection
MRS: De Man Rogosa Sharpe agar
DMEM: Dulbecco’s Modified Eagle Medium
FBS: Fetal Bovine Serum
MTCC: Microbial Type Culture Collection
PBS: Phosphate Buffered Saline
MOI: Multiplicity Of Infection
SS: Salmonella Shigella
DPPH: 1,1-diphenyl-2-picrylhydrazyl
ABTS: 2,2’-azino-bis (3-ethylbenzothiazoline-6-sulfonic acid
PCR: polymerase chain reaction
CFU: colony forming unit
ANI: Average Nucleotide Identity, CRISPR, Clustered Regularly Interspaced Short Palindromic Repeats
SCFA: short-chain fatty acid
PKs: Polyketide Synthases
NRPs: Nonribosomal peptides
RiPP: Ribosomally synthesized and post-translationally modified peptides

## Authors’ Contributions

SP performed all experiments, including analysis, and drafted the manuscript. BP reviewed and edited the manuscript. ARC conceived the study, designed the approach, evaluated the data, and was involved in manuscript preparation. Finally, all authors reviewed and edited the manuscript.

## Ethical statements

The author(s) state that there is no conflict of interest.

## Acknowledgments

This study was supported by the intramural funding provided by IIT Bhubaneswar. SP receives an institutional fellowship from IIT Bhubaneswar. BP is supported by the endowment grant to IIT Bhubaneswar from the Dr. Dash Foundation, USA. We are grateful to Dr. Nrisingha Dey, ILS Bhubaneswar, for providing us Gram-positive pathogens [Methicillin-resistant *Staphylococcus aureus* ATCC 700699 (MRSA) and *Staphylococcus aureus* ATCC 25923] used in this study. All the Gram-negative clinical isolates [*Escherichia coli* (ETEC) BCH 04067*, Vibrio cholerae* BCH 09616*, Shigella Flexneri* BCH 06745] were generous gifts from Dr. Asish Kumar Mukhopadhyay, NICED Kolkata. We are thankful to Dr. Tushar Kant Beuria and his group, ILS Bhubaneswar, for providing technical guidance for the biofilm assays. We would like to thank Prof. Surajit Mandal, West Bengal University of Animal and Fishery Sciences, Kolkata, for his suggestions on the surface hydrophobicity assay. We would like to convey our thanks to Dr. Chandan Goswami, NISER, Bhubaneswar, for providing the confocal microscopy facility.

## Data availability

The whole genome data of *Ligilactobacillus salivarius LS-ARS2* is in NCBI repository, and can be accessed under BioProject: PRJNA1024881 (https://www.ncbi.nlm.nih.gov/sra/?term=PRJNA1024881 OR https://www.ncbi.nlm.nih.gov/search/all/?term=PRJNA1024881).

---

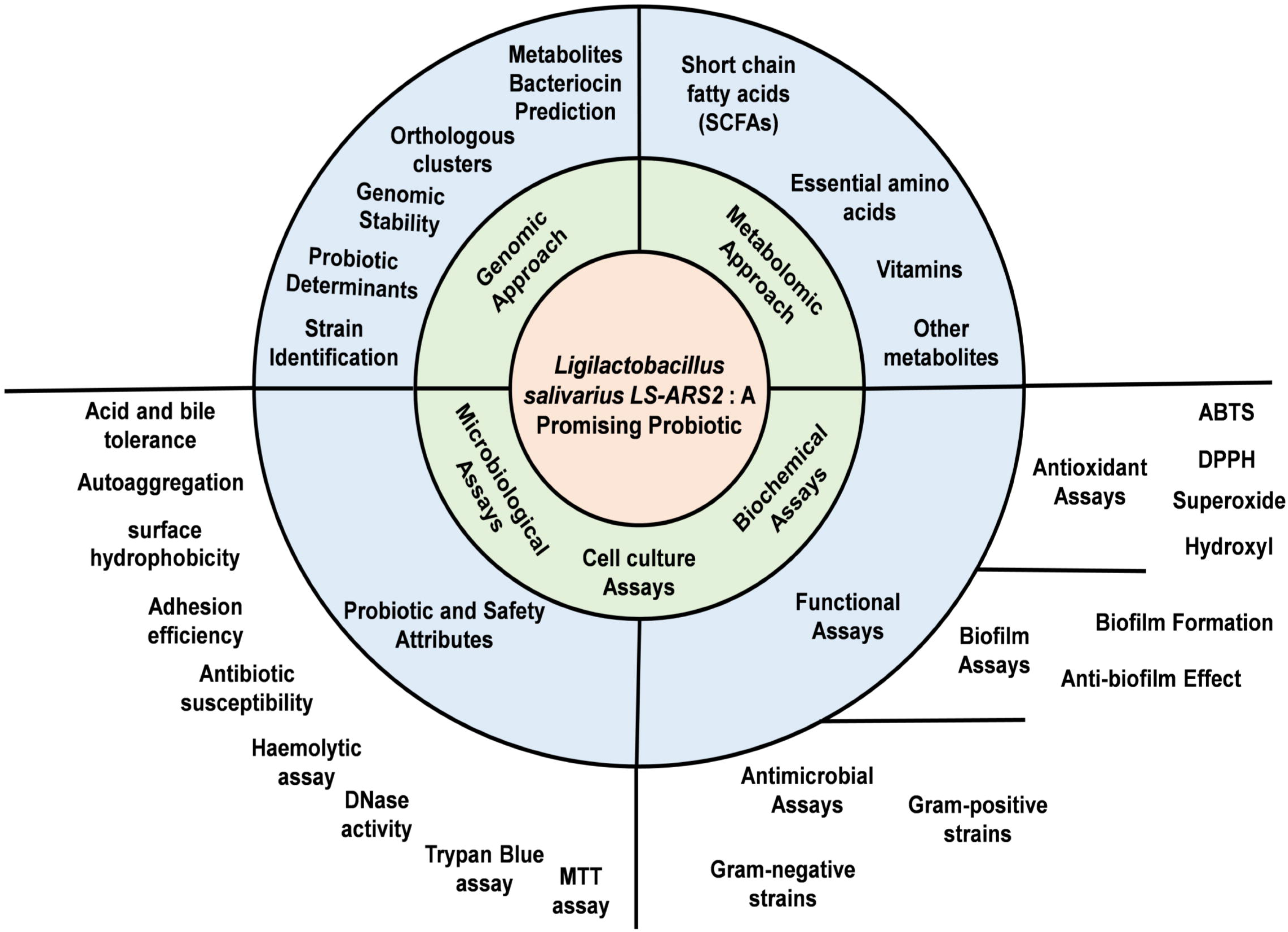

## References

Abouloifa, H., Rokni, Y., Bellaouchi, R., Ghabbour, N., Karboune, S., Brasca, M. et al. (2020). Characterization of probiotic properties of antifungal *Lactobacillus* strains isolated from traditional fermenting green olives. Probiotics Antimicrob. Proteins 12, 683–696. doi: 10.1007/s12602-019-09543-8

Arndt, D., Grant, J.R., Marcu, A., Sajed, T., Pon, A., Liang, Y. et al. (2016). PHASTER: a better, faster version of the PHAST phage search tool. Nucleic Acids Res. 44, W16–W21. doi: 10.1093/nar/gkw387

Arumugam, M.K., Paal, M.C., Donohue Jr, T.M., Ganesan, M., Osna, N.A., Kharbanda, K.K. (2021). Beneficial effects of betaine: A comprehensive review. Biology 10, 456. doi: 10.3390/biology10060456

Ayala, D.I., Cook, P.W., Franco, J.G., Bugarel, M., Kottapalli, K.R., Loneragan, G.H. et al. (2019). A systematic approach to identify and characterize the effectiveness and safety of novel probiotic strains to control foodborne pathogens. Front. Microbiol. 10, 1108. doi: 10.3389/fmicb.2019.01108

Ayed, L., M’hir, S., Nuzzolese, D., Di Cagno, R., Filannino, P. (2024). Harnessing the Health and Techno-Functional Potential of Lactic Acid Bacteria: A Comprehensive Review. Foods 13, 1538. doi: 10.3390/foods13101538

Baldassarri, L., Creti, R., Recchia, S., Imperi, M., Facinelli, B., Giovanetti, E. et al. (2006). Therapeutic failures of antibiotics used to treat macrolide-susceptible *Streptococcus pyogenes* infections may be due to biofilm formation. J. Clin. Microbiol. 44, 2721–2727. doi: 10.1128/jcm.00512-06

Barrett, E., Hayes, M., O’Connor, P., Gardiner, G., Fitzgerald, G.F., Stanton, C. et al. (2007). Salivaricin P, one of a family of two-component antilisterial bacteriocins produced by intestinal isolates of *Lactobacillus salivarius*. Appl. Environ. Microbiol. 73, 3719–3723. doi: 10.1128/AEM.00666-06

Barry, H. (1991). Reactive oxygen species in living systems. Am J Med 91, S14. doi: 10.1016/0002-9343(91)90279-7

Blin, K., Shaw, S., Kloosterman, A.M., Charlop Powers, Z., Van Wezel, G.P., Medema, M.H. et al. (2021). antiSMASH 6.0: improving cluster detection and comparison capabilities. Nucleic Acids Res. 49, W29–W35. doi: 10.1093/nar/gkab335

Bortolaia, V., Kaas, R.S., Ruppe, E., Roberts, M.C., Schwarz, S., Cattoir, V. et al. (2020). ResFinder 4.0 for predictions of phenotypes from genotypes. J. Antimicrob. Chemother. 75, 3491–3500. doi: 10.1093/jac/dkaa345

Bs, S., Thankappan, B., Mahendran, R., Muthusamy, G., Femil Selta, D.R., Angayarkanni, J. (2021). Evaluation of GABA production and probiotic activities of *Enterococcus faecium* BS5. Probiotics Antimicrob. Proteins 13, 993–1004. doi: 10.1007/s12602-021-09759-7

Campana, R., van Hemert, S., Baffone, W. (2017). Strain-specific probiotic properties of lactic acid bacteria and their interference with human intestinal pathogens invasion. Gut Pathog. 9, 1–12. doi: 10.1186/s13099-017-0162-4

Chen, C.C., Lai, C.C., Huang, H.L., Huang, W.Y., Toh, H.S., Weng, T.C. et al. (2019). Antimicrobial activity of *Lactobacillus* species against carbapenem-resistant *Enterobacteriaceae*. Front. Microbiol. 10, 789. doi: 10.3389/fmicb.2019.00789

Ciofu, O., Moser, C., Jensen, P.Ø., Høiby, N. (2022). Tolerance and resistance of microbial biofilms. Nat. Rev. Microbiol. 20, 621–635. doi: 10.1038/s41579-022-00682-4

CLSI, C.L.S.I. (2020). Performance standards for antimicrobial susceptibility testing. 30th edition.

Dash, J., Sethi, M., Deb, S., Parida, D., Kar, S., Mahapatra, S. et al. (2023). Biochemical, functional and genomic characterization of a new probiotic *Ligilactobacillus salivarius* F14 from the gut of tribes of Odisha. World J. Microbiol. Biotechnol. 39, 171. doi: 10.1007/s11274-023-03626-z

de Jong, A., van Hijum, S.A., Bijlsma, J.J., Kok, J., Kuipers, O.P. (2006). BAGEL: a web-based bacteriocin genome mining tool. Nucleic Acids Res. 34, W273–W279. doi: 10.1093/nar/gkl237

Deghorain, M., Goffin, P., Fontaine, L., Mainardi, J.L., Daniel, R., Errington, J. et al. (2007). Selectivity for D-lactate incorporation into the peptidoglycan precursors of *Lactobacillus plantarum*: role of Aad, a VanX-like D-alanyl-D-alanine dipeptidase. J Bacteriol. 189, 4332–4337. doi: 10.1128/jb.01829-06

Dietzsch, S., Braesigk, A., Seidel, C., Remmele, J., Kitzing, R., Schlender, T. et al. (2021). Types of deviation and review criteria in pretreatment central quality control of tumor bed boost in medulloblastoma—an analysis of the German Radiotherapy Quality Control Panel in the SIOP PNET5 MB trial. Strahlenther Onkol 198, 282–290. doi: 10.1007/s00066-021-01822-0

Donia, M.S., Fischbach, M.A. (2015). Small molecules from the human microbiota. Science 349, 1254766. doi: 10.1126/science.1254766

Fonseca, H.C., de Sousa Melo, D., Ramos, C.L., Dias, D.R., Schwan, R.F. (2021). Probiotic properties of *lactobacilli* and their ability to inhibit the adhesion of *enteropathogenic* bacteria to Caco-2 and HT-29 cells. Probiotics Antimicrob. Proteins 13, 102–112. doi: 10.1007/s12602-020-09659-2

Furman, D., Campisi, J., Verdin, E., Carrera Bastos, P., Targ, S., Franceschi, C. et al. (2019). Chronic inflammation in the etiology of disease across the life span. Nat. Med. 25, 1822–1832. doi: 10.1038/s41591-019-0675-0

Gao, D., Gao, Z., Zhu, G. (2013). Antioxidant effects of *Lactobacillus plantarum* via activation of transcription factor Nrf2. Food Funct. 4, 982–989. doi: 10.1039/C3FO30316K

Gao, J., Sadiq, F.A., Zheng, Y., Zhao, J., He, G., Sang, Y. (2022). Biofilm-based delivery approaches and specific enrichment strategies of probiotics in the human gut. Gut Microbes 14, 2126274. doi: 10.1080/19490976.2022.2126274

Gasperi, V., Sibilano, M., Savini, I., Catani, M.V. (2019). Niacin in the central nervous system: an update of biological aspects and clinical applications. Int. J. Mol. Sci. 20, 974. doi: 10.3390/ijms20040974

Grissa, I., Vergnaud, G., Pourcel, C. (2007). CRISPRFinder: a web tool to identify clustered regularly interspaced short palindromic repeats. Nucleic Acids Res. 35, W52–W57. doi: 10.1093/nar/gkm360

Gu, S., Zhao, Z., Yao, Y., Li, J., Tian, C. (2022). Designing and constructing a novel artificial pathway for malonic acid production biologically. Front. Bioeng. Biotechnol. 9, 820507. doi: 10.3389/fbioe.2021.820507

Guerrero Sanchez, M., Passot, S., Campoy, S., Olivares, M., Fonseca, F. (2022). *Ligilactobacillus salivarius* functionalities, applications, and manufacturing challenges. Appl. Microbiol. Biotechnol. 106, 57–80. doi: 10.1007/s00253-021-11694-0

Hakansson, A., Molin, G. (2011). Gut microbiota and inflammation. Nutrients 3, 637–682. doi: 10.3390/nu3060637

Horn, H.J., Holland, E., Hazleton, L. (1957). Food Additives, Safety of Adipic Acid as Compared with Citric and Tartaric Acid. J. Agric. Food Chem. 5, 759–762. doi: 10.1021/jf60080a007

Hsiung, R.T., Fang, W.T., LePage, B.A., Hsu, S.A., Hsu, C.H., Chou, J.Y. (2021). In vitro properties of potential probiotic indigenous yeasts originating from fermented food and beverages in Taiwan. Probiotics Antimicrob. Proteins 13, 113–124. doi: 10.1007/s12602-020-09661-8

Jia, B., Raphenya, A.R., Alcock, B., Waglechner, N., Guo, P., Tsang, K.K. et al. (2016). CARD 2017: expansion and model-centric curation of the comprehensive antibiotic resistance database. Nucleic Acids Res. 45, D566–D573. doi: 10.1093/nar/gkw1004

Jiang, Y.H., Yang, R.S., Lin, Y.C., Xin, W.G., Zhou, H.Y., Wang, F. et al. (2023). Assessment of the safety and probiotic characteristics of *Lactobacillus salivarius* CGMCC20700 based on whole-genome sequencing and phenotypic analysis. Front. Microbiol. 14, 1120263. doi: 10.3389/fmicb.2023.1120263

Jm, M. (1985). Oxygen-derived free radicals in postischemic tissue injury. N Engl J Med 312, 159–163. doi: 10.1056/NEJM198501173120305

Kandasamy, S., Lee, K.H., Yoo, J., Yun, J., Kang, H.B., Kim, J.E. et al. (2024). Whole genome sequencing of *Lacticaseibacillus casei* KACC92338 strain with strong antioxidant activity, reveals genes and gene clusters of probiotic and antimicrobial potential. Front. Microbiol. 15, 1458221. doi: 10.3389/fmicb.2024.1458221

Katsuyama, Y., Ohnishi, Y. (2012) Type III polyketide synthases in microorganisms. Methods Enzymol. 515, 359–377. doi: 10.1016/B978-0-12-394290-6.00017-3

Kaushik, A., Kest, H., Sood, M., Steussy, B.W., Thieman, C., Gupta, S. (2024). Biofilm Producing Methicillin-Resistant *Staphylococcus aureus* (MRSA) Infections in Humans: Clinical Implications and Management. Pathog. 13, 76. doi: 10.3390/pathogens13010076

Kavanova, K., Kostovova, I., Moravkova, M., Kubasova, T., Babak, V., Crhanova, M. (2024). Comparative Genome Analysis and Characterization of the Probiotic Properties of Lactic Acid Bacteria Isolated from the Gastrointestinal Tract of Wild Boars in the Czech Republic. Probiotics Antimicrob. Proteins, 1-19. doi: 10.1007/s12602-024-10259-7

Kim, S., Lee, J.Y., Jeong, Y., Kang, C.H. (2022). Antioxidant activity and probiotic properties of lactic acid bacteria. Fermentation 8, 29. doi: 10.3390/fermentation8010029

Lebeer, S., Vanderleyden, J., De Keersmaecker, S.C. (2008). Genes and molecules of *lactobacilli* supporting probiotic action. *Microbiol*. Mol. Biol. 72, 728–764. doi: 10.1128/mmbr.00017-08

Lee, J.E., Lee, N.K., Paik, H.D. (2021). Antimicrobial and anti-biofilm effects of probiotic *Lactobacillus plantarum* KU200656 isolated from kimchi. Food Sci. Biotechnol. 30, 97–106. doi: 10.1007/s10068-020-00837-0

Lei, Y., Oshima, T., Ogasawara, N., Ishikawa, S. (2013). Functional analysis of the protein Veg, which stimulates biofilm formation in *Bacillus subtilis*. J Bacteriol. 195, 1697–1705. doi: 10.1128/jb.02201-12

Li, H., Durbin, R. (2009). Fast and accurate short read alignment with Burrows–Wheeler transform. bioinformatics 25, 1754–1760. doi: 10.1093/bioinformatics/btp324

Liang, X., Dai, N., Yang, F., Zhu, H., Zhang, G., Wang, Y. (2024). Molecular identification and safety assessment of the potential probiotic strain *Bacillus paralicheniformis* HMPM220325 isolated from artisanal fruit dairy products. Food Funct. 15, 747–765. doi: 10.1039/D3FO04625G

Magnusson, J., Schnürer, J. (2001). *Lactobacillus coryniformis* subsp. *coryniformis* strain Si3 produces a broad-spectrum proteinaceous antifungal compound. Appl. Environ. Microbiol. 67, 1–5. doi: 10.1128/AEM.67.1.1-5.2001

McFarland, L.V., Evans, C.T., Goldstein, E.J. (2018). Strain-specificity and disease-specificity of probiotic efficacy: a systematic review and meta-analysis. Front. Med. 5, 124. doi: 10.3389/fmed.2018.00124

Naughton, L.M., Romano, S., O’Gara, F., Dobson, A.D. (2017). Identification of secondary metabolite gene clusters in the *Pseudovibrio* genus reveals encouraging biosynthetic potential toward the production of novel bioactive compounds. Front. Microbiol. 8, 1494. doi: 10.3389/fmicb.2017.01494

Navarré, A., Nazareth, T., Luz, C., Meca, G., Escrivá, L. (2024). Characterization of lactic acid bacteria isolated from human breast milk and their bioactive metabolites with potential application as a probiotic food supplement. Food Funct. 15, 8087–8103. doi: 10.1039/D4FO02171A

Oien, D.B., Moskovitz, J. (2019). Genetic regulation of longevity and age-associated diseases through the methionine sulfoxide reductase system. Biochim. Biophys. Acta Mol. Basis Dis. 1865, 1756–1762. doi: 10.1016/j.bbadis.2018.11.016

Pascal Andreu, V., Roel Touris, J., Dodd, D., Fischbach, M.A., Medema, M.H. (2021). The gutSMASH web server: automated identification of primary metabolic gene clusters from the gut microbiota. Nucleic Acids Res. 49, W263–W270. doi: 10.1093/nar/gkab353

Patra, S., Sahu, N., Saxena, S., Pradhan, B., Nayak, S.K., Roychowdhury, A. (2022). Effects of probiotics at the interface of metabolism and immunity to prevent colorectal cancer-associated gut inflammation: A systematic network and meta-analysis with molecular docking studies. Front. Microbiol. 13, 878297. doi: 10.3389/fmicb.2022.878297

Patra, S., Saxena, S., Sahu, N., Pradhan, B., Roychowdhury, A. (2021). Systematic network and meta-analysis on the antiviral mechanisms of probiotics: a preventive and treatment strategy to mitigate SARS-CoV-2 infection. Probiotics Antimicrob. 13, 1138–1156. doi: 10.1007/s12602-021-09748-w

Pedroza Matute, S., Iyavoo, S. (2023). Exploring the gut microbiota: lifestyle choices, disease associations, and personal genomics. Front. Nutr. 10, 1225120. doi: 10.3389/fnut.2023.1225120

Pei, Z., Sadiq, F.A., Han, X., Zhao, J., Zhang, H., Ross, R.P. et al. (2021). Comprehensive scanning of prophages in *Lactobacillus*: distribution, diversity, antibiotic resistance genes, and linkages with CRISPR-Cas systems. Msystems 6, e01211–01220. doi: 10.1128/msystems.01211-20

Pereira, J., Castro, M.M., Santos, F., Jesus, A.R., Paiva, A., Oliveira, F. et al. (2022). Selective terpene based therapeutic deep eutectic systems against colorectal cancer. Eur J Pharm Biopharm. 175, 13–26. doi: 10.1016/j.ejpb.2022.04.008

Piqué, N., Berlanga, M., Miñana Galbis, D. (2019). Health benefits of heat-killed (Tyndallized) probiotics: an overview. Int. J. Mol. Sci. 20, 2534. doi: 10.3390/ijms20102534

Pradhan, B., Guha, D., Ray, P., Das, D., Aich, P. (2016). Comparative analysis of the effects of two probiotic bacterial strains on metabolism and innate immunity in the RAW 264.7 murine macrophage cell line. Probiotics Antimicrob. Proteins 8, 73–84. doi: 10.1007/s12602-016-9211-4

Ramalho, J.B., Soares, M.B., Spiazzi, C.C., Bicca, D.F., Soares, V.M., Pereira, J.G. et al. (2019). In vitro probiotic and antioxidant potential of *Lactococcus lactis* subsp. *cremoris* LL95 and its effect in mice behaviour. Nutrients 11, 901. doi: 10.3390/nu11040901

Ren, J., Li, H., Zeng, G., Pang, B., Wang, Q., Wei, J. (2023). Gut microbiome-mediated mechanisms in aging-related diseases: are probiotics ready for prime time? Front. Pharmacol. 14, 1178596. doi: 10.3389/fphar.2023.1178596

Rocchetti, M.T., Russo, P., De Simone, N., Capozzi, V., Spano, G., Fiocco, D. (2024). Immunomodulatory activity on human macrophages by cell-free supernatants to explore the probiotic and postbiotic potential of *Lactiplantibacillus plantarum* strains of plant origin. Probiotics Antimicrob. Proteins 16, 911–926. doi: 10.1007/s12602-023-10084-4

Rosenberg, M., Gutnick, D., Rosenberg, E. (1980). Adherence of bacteria to hydrocarbons: a simple method for measuring cell-surface hydrophobicity. FEMS Microbiol. Lett. 9, 29–33. doi: 10.1111/j.1574-6968.1980.tb05599.x

Rychen, G., Aquilina, G., Azimonti, G., Bampidis, V., de Lourdes Bastos, M., Bories, G. et al. (2018). Guidance on the characterisation of microorganisms used as feed additives or as production organisms. Efsa Journal 16, e05206. doi: 10.2903/j.efsa.2018.5206

Salas-Jara, M.J., Ilabaca, A., Vega, M., García, A. (2016). Biofilm forming *Lactobacillus*: new challenges for the development of probiotics. Microorganisms 4, 35. doi: 10.3390/microorganisms4030035

Saxami, G., Kerezoudi, E.N., Eliopoulos, C., Arapoglou, D., Kyriacou, A. (2023). The gut–organ axis within the human body: gut dysbiosis and the role of prebiotics. Life 13, 2023. doi: 10.3390/life13102023

Seemann, T. (2014). Prokka: rapid prokaryotic genome annotation. Bioinformatics 30, 2068–2069. doi: 10.1093/bioinformatics/btu153

Shrestha, P., Karmacharya, J., Han, S.R., Lee, J.H., Oh, T.J. (2024). Elucidation of bacterial trehalose-degrading trehalase and trehalose phosphorylase: physiological significance and its potential applications. Glycobiol 34, cwad084. doi: 10.1093/glycob/cwad084

Stach, K., Stach, W., Augoff, K. (2021). Vitamin B6 in health and disease. Nutrients 13, 3229. doi: 10.3390/nu13093229

Tamura, K., Stecher, G., Kumar, S. (2021). MEGA11: molecular evolutionary genetics analysis version 11. Mol. Biol. Evol. 38, 3022–3027. doi: 10.1093/molbev/msab120

Tenea, G.N., Ascanta, P. (2022). Bioprospecting of ribosomally synthesized and post-translationally modified peptides through genome characterization of a novel probiotic *Lactiplantibacillus plantarum* UTNGt21A strain: A promising natural antimicrobials factory. Front. Microbiol. 13, 868025. doi: 10.3389/fmicb.2022.868025

Tenea, G.N., Ortega, C. (2021). Genome characterization of *Lactiplantibacillus plantarum* strain UTNGt2 originated from *Theobroma grandiflorum* (white cacao) of Ecuadorian Amazon: Antimicrobial peptides from safety to potential applications. Antibiotics 10, 383. doi: 10.3390/antibiotics10040383

Timmerman, H., Veldman, A., Van den Elsen, E., Rombouts, F., Beynen, A. (2006). Mortality and growth performance of broilers given drinking water supplemented with chicken-specific probiotics. Poult. Sci. 85, 1383–1388. doi: 10.1093/ps/85.8.1383

Wernerman, J., Hammarqvist, F., Vinnars, E. (1990). α-Ketoglutarate and postoperative muscle catabolism. Lancet 335, 701–703. doi: 10.1016/0140-6736(90)90811-I

Wong, A., Ngu, D.Y.S., Dan, L.A., Ooi, A., Lim, R.L.H. (2015). Detection of antibiotic resistance in probiotics of dietary supplements. Nutr. J. 14, 1–6. doi: 10.1186/s12937-015-0084-2

Xiong, L., Ni, X., Niu, L., Zhou, Y., Wang, Q., Khalique, A. et al. (2019). Isolation and preliminary screening of a *Weissella confusa* strain from giant panda (*Ailuropoda melanoleuca*). Probiotics Antimicrob. Proteins 11, 535–544. doi: 10.1007/s12602-018-9402-2

Xiong, Z.Q., Kong, L.H., Wang, G.Q., Xia, Y.J., Zhang, H., Yin, B.X. et al. (2018). Functional analysis and heterologous expression of bifunctional glutathione synthetase from *Lactobacillus*. J. dairy sci. 101, 6937–6945. doi: 10.3168/jds.2017-14142

Yang, Y., Song, X., Wang, G., Xia, Y., Xiong, Z., Ai, L. (2024). Understanding *Ligilactobacillus salivarius* from Probiotic Properties to Omics Technology: A Review. Foods 13, 895. doi: 10.3390/foods13060895

Zankari, E., Hasman, H., Cosentino, S., Vestergaard, M., Rasmussen, S., Lund, O. et al. (2012). Identification of acquired antimicrobial resistance genes. J. Antimicrob. Chemother. 67, 2640–2644. doi: 10.1093/jac/dks261

Zhang, W., Ji, H., Zhang, D., Liu, H., Wang, S., Wang, J. et al. (2018). Complete genome sequencing of *Lactobacillus plantarum* ZLP001, a potential probiotic that enhances intestinal epithelial barrier function and defense against pathogens in pigs. Front. physiol. 9, 1689. doi: 10.3389/fphys.2018.01689

Zhao, P., Tian, P. (2021). Biosynthesis pathways and strategies for improving 3-hydroxypropionic acid production in bacteria. World J. Microbiol. Biotechnol. 37, 117. doi: 10.1007/s11274-021-03091-6

Zheng, J., Ge, Q., Yan, Y., Zhang, X., Huang, L., Yin, Y. (2023). dbCAN3: automated carbohydrate-active enzyme and substrate annotation. Nucleic Acids Res. 51, W115–W121. doi: 10.1093/nar/gkad328

