## Supplementary material for "Complete Genome Sequence, Metabolic Profiling and Functional Studies reveal *Ligilactobacillus salivarius LS-ARS2* is a Promising Biofilm-forming Probiotic with Significant Antioxidant, Antibacterial, and Antibiofilm Potential": pdf attached

### Supplementary Figures

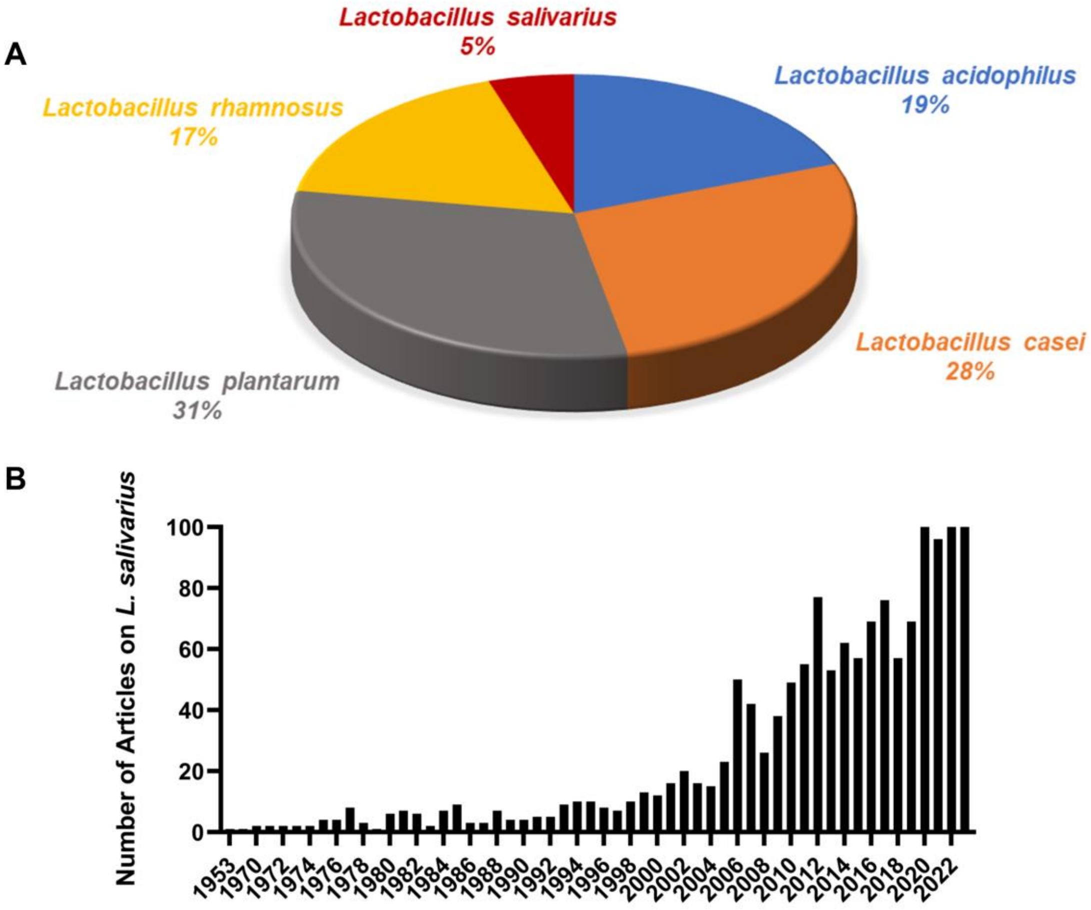


**Supplementary Figure S1 Updated scenario of research on *Lactobacillus salivarius* strains. (A)** Research trend showing that compared to other LAB, available studies on *Lactobacillus salivarius* strains are limited. **(B)** Number of research article published on *L. salivarius* is consistently increased showing the promising attributes of *L. salivarius.*

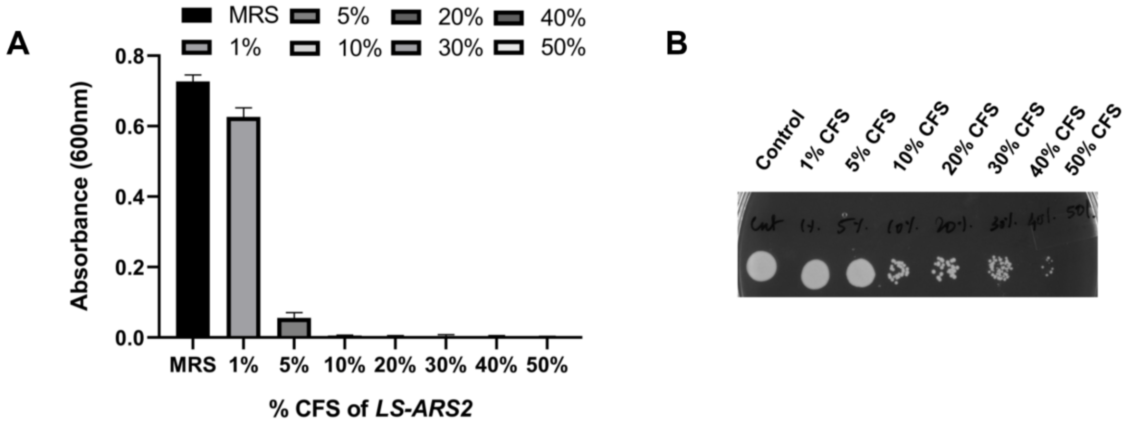


**Supplementary Figure S2 Determination of minimum inhibitory percentage (MIP) of *LS-ARS2*-derived CFS for Methicillin-resistant *Staphylococcus aureus* (MRSA) (A) To determine the MIP,** MRSA was grown in different percentages of *LS-ARS2* CFS. **(B) To determine the viability of MRSA**, from each well of pathogen cultured in *LS-ARS2* CFS, spotting was performed and checked whether the effect was bactericidal or bacteriostatic. All the data were represented as mean ± SD of three replicates.

### Supplementary Tables

**Supplementary Table S1 Overall assembly summary**

| Sample name | Raw data | Data after AT (Gb) | Read quality before AT | Read quality after AT | GC% before AT | GC% after AT | Coverage (>=30x) | Depth |
|---|---|---|---|---|---|---|---|---|
| ARS-2 | 1.22 | 0.85 | 37.75 | 37.74 | 35.5 | 33.35 | 90.12% | 290.81 |

Data after AT (Gb): Data after Adapter Trimming

Read quality before AT: Read quality before Adapter Trimming

Read quality after AT: Read quality after Adapter Trimming

GC% before AT: GC% before Adapter Trimming

GC% after AT: GC% after Adapter Trimming

Coverage (%): The percentage of mapped sited (>= 1x)

Depth: Average mapping depth

**Supplementary Table S2 Functions of the relevant genes obtained from the *LS-ARS2* genome**

| Preferred_name | Function |
|---|---|
| clpC | Part of a stress-induced multi-chaperone system |
| yhcA | ABC transporter, ATP-binding protein |
| lmrA | ABC transporter, ATP-binding protein |
| yfiC | ABC transporter |
| ybhF_2 | AAA domain, putative AbiEii toxin, Type IV TA system |
| ybhR | ABC transporter |
| VPA1512 | Bacterial extracellular solute-binding proteins, family 3 |
| **veg** | **Biofilm formation stimulator VEG** |
| argH | argininosuccinate lyase |
| glnP | ABC transporter |
| uspA | Universal stress protein family |
| ytmP | Choline/ethanolamine kinase |
| ecsB | ABC transporter |
| comA | ABC-type bacteriocin lantibiotic exporters |
| grpE | Participates in the response to hyperosmotic and heat shock |
| dnaK | Heat shock 70 kDa protein |
| glnP | ABC transporter |
| glnQ | ABC transporter, ATP-binding protein |
| atpB | it plays a direct role in the translocation of protons across the membrane |
| clpB | Part of a stress-induced multi-chaperone system |
| natB | ABC-type Na efflux pump, permease component |
| **dltD** | **Protein involved in D-alanine esterification of lipoteichoic acid and wall teichoic acid (D-alanine transfer protein)** |
| **dltC** | **Carrier protein involved in the D-alanylation of lipoteichoic acid (LTA).** |
| **dltB** | **MBOAT, membrane-bound O-acyltransferase family** |

| | |
|---|---|
| **dltA** | **Catalyzes the first step in the D-alanylation of lipoteichoic acid (LTA)** |
| **dltX** | **D-Ala-teichoic acid biosynthesis protein** |
| mntH | H (+)-stimulated, divalent metal cation uptake system |
| nrdG | Activation of anaerobic ribonucleoside-triphosphate reductase under anaerobic conditions |
| glnQ | ABC transporter, ATP-binding protein |
| tcyB | ABC transporter |
| yjcE | Sodium proton antiporter |
| hrtB | ABC transporter permease |
| devA | ABC transporter, ATP-binding protein |
| **luxS** | **Involved in the synthesis of autoinducer 2 (AI-2) and quorum sensing** |
| uspA | universal stress protein |
| cydD | ABC transporter transmembrane region |
| cydD | ABC transporter, CydDC cysteine exporter (CydDC-E) family, permease ATP-binding protein CydD |
| aatB | ABC transporter substrate-binding protein |
| glnQ | ABC transporter, ATP-binding protein |
| glnP | ABC transporter permease |
| fhuC | ABC transporter |
| znuB | ABC 3 transport family |
| glnPH2 | ABC transporter permease |
| glnQ | ABC transporter, ATP-binding protein |
| cas3 | CRISPR-associated helicase cas3 |
| casB | CRISPR-associated protein Cse2 (CRISPR_cse2) |
| casD | CRISPR-associated protein (Cas_Cas5) |
| casE | CRISPR_assoc |
| cas1 | Provides protection against mobile genetic elements (viruses, transposable elements and conjugative plasmids) |
| cas2 | CRISPR-associated protein (Cas_Cas2CT1978) |
| metI | ABC transporter permease |
| metN | Part of the ABC transporter complex MetNIQ involved in methionine import |
| metI | ABC transporter permease |
| usp6 | universal stress protein |

| | |
|---|---|
| pflA | Activation of pyruvate formate-lyase under anaerobic conditions |
| potD | ABC transporter |
| potB | ABC transporter permease |
| **tpx** | **Thiol-specific peroxidase that catalyzes reduction of hydrogen peroxide and organic hydroperoxides to water and alcohols, respectively** |
| **yfeX** | **Peroxidase** |
| **trxA** | **Belongs to the thioredoxin family** |
| **trxB** | **Belongs to the class-II pyridine nucleotide-disulfide oxidoreductase family** |
| **yjbH** | **Thioredoxin** |
| **gshF** | **Belongs to the glutamate--cysteine ligase type 1 family** |
| **ndh** | **NADH dehydrogenase** |
| **npr** | **Pyridine nucleotide-disulphide oxidoreductase, dimerisation domain** |
| **nrdH** | **Glutaredoxin** |
| **msrA** | **Catalyzes the reversible oxidation-reduction of methionine sulfoxide in proteins to methionine and protects proteins from oxidation** |
| **msrB** | **peptide methionine sulfoxide reductase** |
| **pts13C** | **The phosphoenolpyruvate-dependent sugar phosphotransferase system (PTS), a major carbohydrate active - transport system** |
| **hprK** | **Participate in sugar transport by sugar phosphotransferase system (PTS)** |
| **epsL** | **Bacterial sugar transferase** |
| **mtlF** | **Catalyzes the phosphorylation of incoming sugar substrates concomitant with their translocation across the cell membrane** |
| **ptsI** | **Component of the phosphoenolpyruvate-dependent sugar phosphotransferase system (sugar PTS)** |
| **uxuT** | **MFS/sugar transport protein** |
| **glcU** | **sugar transport** |
| **rfbP** | **Bacterial sugar transferase** |
| **manL** | **PTS system sorbose subfamily IIB component** |
| **dhaM** | **PTS system fructose IIA component** |
| **srlB** | **PTS system glucitol/sorbitol-specific IIA component** |
| **srlA** | **PTS system enzyme II sorbitol-specific factor** |

| | |
|---|---|
| **mtlA** | **PTS system, Lactose/Cellobiose specific IIB subunit** |
| **manY** | **PTS system sorbose-specific iic component** |
| **manL** | **PTS system sorbose subfamily IIB component** |
| **comA** | **ABC-type bacteriocin lantibiotic exporters** |

**Supplementary Table S3 Predicted prophage regions within the genome of *LS-ARS2***

| Contig | Region | Region Length (kb) | Completeness | Score | Total Protein | Region Position | Most Common Phage | GC% |
|---|---|---|---|---|---|---|---|---|
| 28 | 1 | 8.4 | Incomplete | 10 | 8 | 352-8848 | PHAGE_Prochl_P_SSM2_NC_006883(4) | 33.73 |
| 48 | 1 | 10 | Incomplete | 10 | 12 | 27884-37902 | PHAGE_Entero_phiEF24C_NC_00904(3) | 31.48 |
| | 2 | 7.9 | Incomplete | 10 | 8 | 41090-49082 | PHAGE_Cellul_phi38:1_NC_021796(1) | 32.50 |
| 77 | 1 | 9.1 | Incomplete | 10 | 8 | 49876-59038 | PHAGE_Bacill_G_NC_023719(2) | 34.30 |

**Supplementary Table S4 CRISPR-Cas arrays in the *LS-ARS2* genome**

| Element | CRISPR Id / Cas Type | Start | End | Spacer / Gene | Repeat consensus / cas genes |
|---|---|---|---|---|---|
| Cas cluster | CAS | 10790 | 20037 | 7 | cas2_TypeIE, cas3_TypeI, cas3_TypeI, cas5_TypeIE, cas6_TypeIE, cas7_TypeIE, cse2_TypeIE |
| Cas cluster | CAS-TypeIE | 15249 | 20037 | 6 | cas1_TypeIE, cas2_TypeIE, cas5_TypeIE, cas6_TypeIE, cas7_TypeIE, cse2_TypeIE |
| Cas cluster | CAS | 23110 | 25140 | 2 | cas3_TypeI, cas3_TypeI |
| Cas cluster | CAS | 7729 | 9051 | 2 | cas3_TypeI, cas3_TypeI |
| CRISPR | JVAF01000101_1_1 | 56 | 146 | 1 | AAAGTAATACCAATCGTTACCACCTTG |
| CRISPR | JVAF01000101_1_2 | 245 | 394 | 2 | AAAGTAATACCAATCGTTACCACCTTG |
| CRISPR | JVAF01000103_1_1 | 19887 | 19978 | 1 | TTCAATCCAACAAGTGGACGTATGCAAAA |
| Cas cluster | CAS | 37512 | 49026 | 3 | cas4_TypeI-II, cas3_TypeI, cas3_TypeI |

**Supplementary Table S5 Antibiotic resistance gene family, drug class and resistance mechanism of *LS-ARS2***

| RGI criteria | ARO term | Detection criteria | AMR gene family | Drug class | Resistance mechanism | % Identity of Matching region | % Length of Reference Sequence |
|---|---|---|---|---|---|---|---|
| Strict | vanT gene in vanG cluster | protein homolog model | glycopeptide resistance gene cluster, vanT | glycopeptide antibiotic | antibiotic target alteration | 35.52 | 52.11 |

**Supplementary Table S6 Identified genes within the T3PKS secondary metabolite biosynthetic gene clusters with antiSMASH**

| Name | Category | Function | *Score | **E-value |
|---|---|---|---|---|
| T3PKS: Chal_sti_synt_N SMCOG1043: hydroxymethylglutaryl-CoA synthase | Biosynthetic | Cholesterol biosynthesis | 527.6 | 2.9e-160 |
| SMCOG1063: argininosuccinate lyase/adenylosuccinate lyase | Biosynthetic additional | Amino acid biosynthesis | 226.7 | 9.4e-69 |
| ATP-grasp | | Bacterial cell wall biosynthesis | | |
| SMCOG1182: Polyprenyl synthetase | | Secondary metabolites (isoprenoids) | 238.9 | 1.1e-72 |
| SMCOG1008: response regulator | Regulatory | Stress adaption of bacteria | 205.9 | 9.7e-63 |
| SMCOG1003: sensor histidine kinase | | | 198.8 | 3.4e-60 |

*Score and **E-value: Indicates the quality of match between the *LS-ARS2* gene sequences with the reference sequences in the database
